# De novo design of light-controlled conformational protein switches

**DOI:** 10.64898/2026.10.26.761880

**Authors:** Alexander T. Hilditch, Nicolas M. Goldbach, Chenxuan Liu, Luciano A. Abriata, Ann-Sophie Kröll, Friederike C. Reinhardt, Sabine Aschenbrenner, Natalia Gasilova, Sarah Warrelmann, René Rasche, Yassine Faik, Kimali Pilet, Elisabetta Occelli, Sandrine Georgeon, Joseph Schmidt, Laure Menin, Joerg Standfuss, Josef Wachtveitl, Dominik Niopek, Bruno E. Correia

**Author notes:** Equal contribution (Alexander T. Hilditch, Nicolas M. Goldbach).

## Abstract

Light-responsive proteins offer the ability to control biology at an unmatched resolution. Yet, optogenetics is limited to a relatively small number of natural and engineered photoreceptors. The design of light-responsive proteins with native-like complexity is challenging as they must incorporate a light-responsive chromophore, couple chemical changes in the chromophore to structural changes in the protein, and ultimately propagate these changes to downstream effectors. Here, we present the computational design of new light-responsive proteins entirely from the bottom-up. We design *de novo* proteins that accurately bind a retinal chromophore and further design a protein conformational switching motion linked to retinal isomerisation. This switch unlocks a new protein-peptide interaction, creating light-activated heterodimers with an over 30-fold increase in binding upon illumination. Moreover, we show that the designed heterodimers are customisable, with tuneable wavelengths, sensitivities, and specificities. Further, we use the designed heterodimers to control biological function, regulating intracellular protein-protein interactions and cell-signalling receptors. Finally, we use real-time light-activated NMR to demonstrate that they undergo a defined light-activated conformational change. Overall, we computationally design a new class of light-responsive proteins that may open a new optogenetics toolbox with applications in biology.

## Main

Light-responsive proteins have been transformative tools in many domains of biology. Light provides spatial-temporal control with a resolution that is unmatched by either chemical or genetic perturbations. Light-responsive proteins have been used across biology with many applications, from the modulation of cell signalling pathways and control of gene expression^1,2^, to whole-cell neuronal activation^3^. However, despite their broad utility, few light-responsive proteins are available for optogenetics^4^. Natural light-responsive proteins have evolved according to their host’s specific needs, presenting extraordinary levels of specificity, with precise wavelengths, specific chromophores, and light-switching mechanisms that arose from evolutionary optimisation. While powerful in many different ways, their natural origin means that they frequently translate poorly to new contexts, and resist engineering beyond their natural properties. Instead, protein design provides a means for the design of novel proteins that have not been optimised for any particular biological context. Thus, designed light-responsive proteins could offer potentially unmet flexibility, and enable the customisable control of biomolecules using light, creating a valuable optogenetics toolbox for dissecting biological mechanisms and for applications in synthetic biology^5^.

Much of the challenge in designing light-responsive proteins stems from their mechanistic complexity. Light-responsive proteins must incorporate a light-sensitive chromophore, and transduce chemical changes in the chromophore into broader conformational changes in the sensor protein to trigger downstream effects. To-date, engineered light-responsive proteins have borrowed heavily from natural scaffolds and mechanisms^6,7^, or have bypassed the need for a conformational change by using light-responsive moieties directly on a PPI interface^8^. With recent advances in deep learning, protein design promises to expand the toolbox of protein sensors and effectors to sense and control biology^9^. However, proteins that accurately bind small molecules or change conformation are still challenging design targets^10^. The design of proteins with conformational changes requires defining and generating two or more conformations, which are designable, and have a feasible structural transition between them^11^. Major efforts have focused on generating protein conformational ensembles or inverse folding models conditioned for dynamics^12,13^, yet so far their application lags behind the generalisability of single state methods. As a result, few *de novo* designed proteins incorporate stimulus-responsive conformational changes, with notable examples including calcium-controlled dynamic proteins^14^, helical hinge proteins^15^, and protein phosphoswitches^16,17^.

Here, we use computational protein design to create *de novo* light-responsive proteins, designing ligand binding, light-activated conformational changes by coupling protein dynamics to ligand conformational changes, and a *de novo* conformation-specific PPI entirely from the bottom up (Fig. 1a). We create soluble *de novo* proteins that incorporate the light-sensitive small molecule retinal, naturally found in the rhodopsin families of membrane-bound photoreceptors (Fig. 1b). We design a conformational switch linked to retinal isomerisation that unveils a protein-protein interaction (PPI) interface, creating light-activated heterodimers (Fig. 1c). These heterodimers show specific light-response, with controllable interactions and wavelengths. Further, we show that the designed light-activated heterodimers can control cellular processes by activating intracellular protein-protein interactions and cell surface receptors in response to light. Overall, these light-responsive designs mark an important step towards expanding the optogenetics toolbox through the development of entirely novel light sensors with features distinct from those found in nature.

**Fig. 1:**
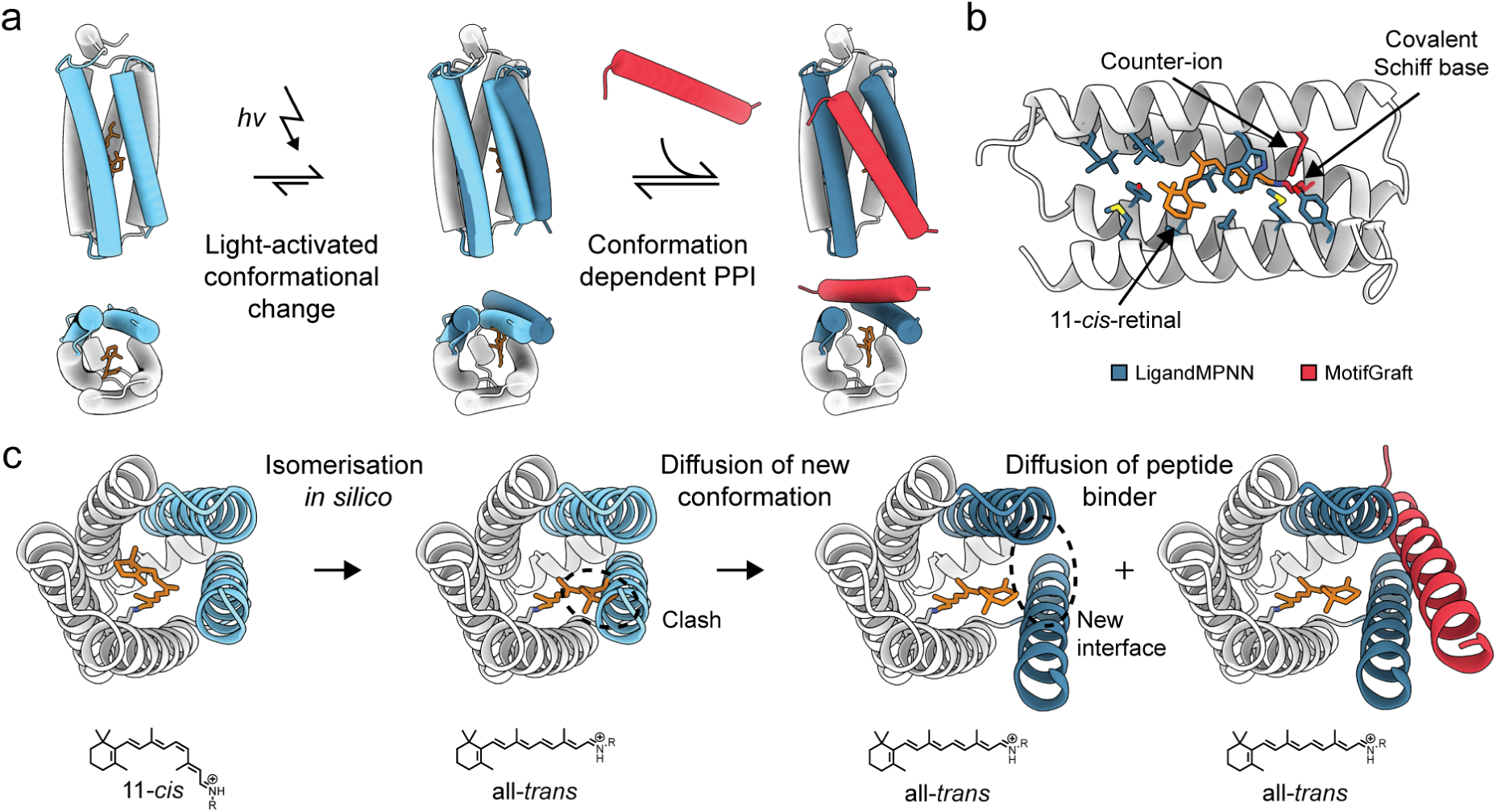
*De novo* design of light-responsive proteins. **a,** Conceptual schematic for the design of light-responsive proteins. Native-like light-responsive proteins must incorporate a light-activated conformational change, and active-state specific function, such as a PPI. **b,** *De novo* retinal binding pockets, with retinal covalently bound via a designed Schiff base linkage. Key residues (lysine and counter-ion) were designed using Rosetta MotifGraft, and the remainder of the pocket using LigandMPNN. **c,** The design of light-activated conformational changes linked to retinal isomerisation. Retinal is modelled in the all-*trans* isomeric state, and RFDiffusionAllAtom used to generate a new conformation. Simultaneously, a new peptide binder is diffused to be complementary to the new conformation.

### *De novo* retinal-binding proteins

In bio-compatible light-responsive systems there are few molecules that could be considered as chromophores for *de novo* proteins. Flavin and phytochrome-based proteins have been used most widely and successfully in soluble light-responsive proteins^18,19^, yet both have limitations. Flavins are largely restricted to blue light activation^20^, while both phytochromes and their bilin chromophores are large molecules, making both their design and use more challenging^21^. Here, we chose to use retinal as a chromophore. Retinal is a uniquely attractive chromophore for the design of light-responsive domains – rhodopsins cover the entire visible spectrum of light^22,23^, and retinal undergoes a *cis* to *trans*, or *trans* to *cis* isomerisation in response to light, resulting in a large, well-characterised conformational change^24^. The limitation is that retinal is not naturally used as a chromophore in soluble proteins, rendering its compatibility with soluble light-responsive proteins unexplored.

We rationalised that by designing topologically simple yet highly designable protein components from the bottom-up, we could build ligand binding, conformational switching, and a *de novo* PPI into the same scaffold. To this end, we designed a range of retinal binding proteins (RBPs) using α-helical bundle (4, 5, and 6-helix bundles)^25^ and β-barrel scaffolds to bind 11-*cis*-retinal^26^. Here, in contrast to previous soluble analogues of rhodopsins or engineered proteins^27–29^, the retinal binding pockets were designed entirely *de novo,* with retinal linked via a Schiff base to a core lysine, supported by a counter-ion and hydrogen bonding network to induce protonation^30^, in a fully *de novo* binding pocket.

As protonated Schiff base formation induces a red shifting in retinal absorbance, designs were screened spectroscopically. Putative hits were then selected for purification and validation. For each of the folds tested we identified designs causing red-shifting of retinal absorbance, with design success rates ranging from 2.5% (β -barrel) to 13.5% (5-helix bundle; Supplementary Figs. 1 & 2). Moreover, analogous to natural rhodopsins, the designs produced varying degrees of red-shifting, with maxima ranging from 440 to 530 nm (unbound retinal – 380 nm; Fig. 2a, Supplementary Fig. 3).

**Fig. 2:**
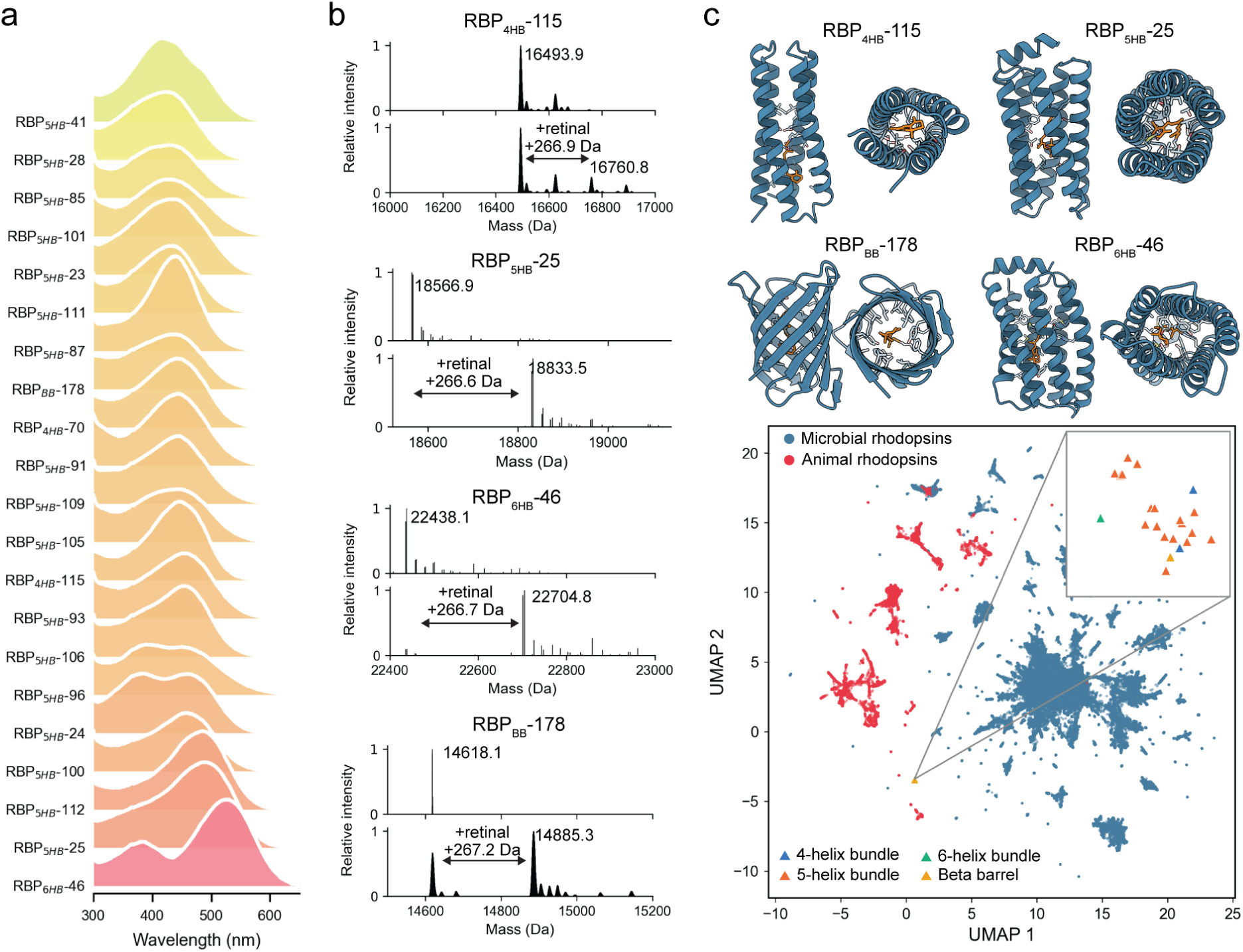
Experimental characterisation of *de novo* designed retinal-binding proteins. **a,** Absorbance spectra of all retinal binding designs made in the study, in the visible range. **b,** Native mass spectra of one protein from each designed fold showing formation of the protein-retinal complex. **c,** AF3 models of the designs measured in **b,** and a representation of sequence embedding space for natural rhodopsins from the UniRef100 database, and the *de novo* RBPs created in this study.

Retinal binding for each of the target folds was validated by mass spectrometry (MS). Native MS of the protein-ligand complexes indicated a modification of +266 Da, corresponding to retinal Schiff base formation (Fig. 2b), while top-down MS fragmentation confirmed the residue of retinal binding (Supplementary Fig. 4). Following, circular dichroism (CD) spectroscopy confirmed that the designs were folded and highly thermostable, with characteristic fingerprints for well-folded α-helical or β-sheet structured proteins (Supplementary Fig. 5).

Overall, these data confirm that *de novo* proteins can be designed to bind retinal using a variety of topologies. To explore this idea further, we asked whether the designed RBPs had explored novel sequence space in comparison to rhodopsins. To this end, we generated sequence embeddings from natural rhodopsins and the *de novo* designed RBPs using the ESM2 protein language model (Fig. 2c)^31^. Representation of these embeddings shows clusters of the microbial and animal rhodopsin families, with little overlap between them, as would be expected from their convergent evolution and structural dissimilarity. Moreover, the *de novo* designed proteins do not cluster with either family, instead forming an isolated population where all of the RBPs are tightly clustered. Overall, this indicates that the designed proteins are indeed exploring novel sequence space even in comparison to natural proteins evolved to perform a similar function (retinal-based photoswitching).

### Computational design of light-activated heterodimers

Having developed *de novo* proteins that bind retinal, we next sought to design sequences which encoded a mechanism of light response. Inspired by natural light-responsive domains, where activation causes backbone conformational changes in the protein, we conceptualised that light activation of retinal and its *cis*-*trans* isomerisation should cause similar backbone-level rearrangements in the *de novo* proteins. This requires the generation of alternative protein conformations, which are accessible from the starting state while still being different enough to have distinct functions.

Therefore, starting from experimentally validated RBPs, we modelled retinal isomerisation from 11-*cis* to all-*trans in silico*. Here, several RBPs showed steric clashes with the modelled all-*trans* configuration, indicating that all-*trans*-retinal is not accommodated in the structure. Then, to generate alternative conformations of the RBPs we used RFDiffusionAllAtom to diffuse the clashing part of the structure, creating new conformational states that are compatible with all-*trans*-retinal^32^. The resulting structural rearrangements opened a binding site between two helices or β-strands, to which we simultaneously diffused a 25-30 amino acid peptide, creating a binding partner for a site that is coupled to the all-*trans*-retinal, effectively encoding the mechanistic steps of a light-activated heterodimer. Overall, this design approach enables the modelling and design of alternative conformations informed by the retinal’s isomerisation in response to light activation. We name this concept <u>S</u>ynthetic <u>P</u>hoto-<u>A</u>ctivated <u>R</u>etinal-based <u>C</u>oupling (SPARC; Fig. 1c). Testing all our RBPs as putative ground (dark) states in SPARC, we found 5-helix RBPs to be ideal as they pack tightly around the 11-*cis*-retinal, and the modelled alternative conformation could be sufficiently distinct from the ground state. In effect, as the newly diffused peptide was always α-helical, this had the result of turning a monomeric 5-helix bundle in the dark state into a heterodimeric 6-helix bundle upon illumination. We then designed amino acid sequences using either ProteinMPNN^33^, or LigandMPNN^34^, with tied positions^15^ across the two conformations of the 5-helix bundle (SPARC_prot_) while designing the peptide (SPARC_pep_) only in its bound state. We predicted the designed sequences with AlphaFold2 (AF2)^35^ and AlphaFold3 (AF3)^36^ in the dark (peptide absent) and light (peptide present) conditions and selected sequences for which the AF3 predictions matched the modelled conformations and expected retinal isomer.

In total, 92 heterodimers were selected for experimental testing. The designed pairs were initially co-expressed in *E. coli,* each component fused to one half of a split-luciferase (Fig. 3a)^37^. The cells were lysed, the lysate incubated with 11-*cis*-retinal, and the luminescence measured before and after illumination with blue light. Of the 92 initial designs, one design (SPARC-39), showed a consistent increase in luminescence after illumination (Supplementary Fig. 6). Purification of the two components (SPARC-39_prot_ and SPARC-39_pep_) verified that the design produced a modest 1.5-fold increase in luminescence after illumination (Fig. 3b). Measurement without 11-*cis*-retinal confirmed that this response was specific to the presence of retinal, and we validated that the designed switch had retained accurate retinal-binding by measurement of the red-shifting, native MS, and top-down MS fragmentation (Fig. 3c, d, Supplementary Fig. 7). Moreover, mutation of the core lysine for retinal Schiff base formation (K76A) completely abrogated the light-response. In addition, measurement of the parent single-state design RBP_5HB_-87 with SPARC-39_pep_ confirmed that the light response was specific to the multi-state design (Supplementary Fig. 8).

**Fig. 3:**
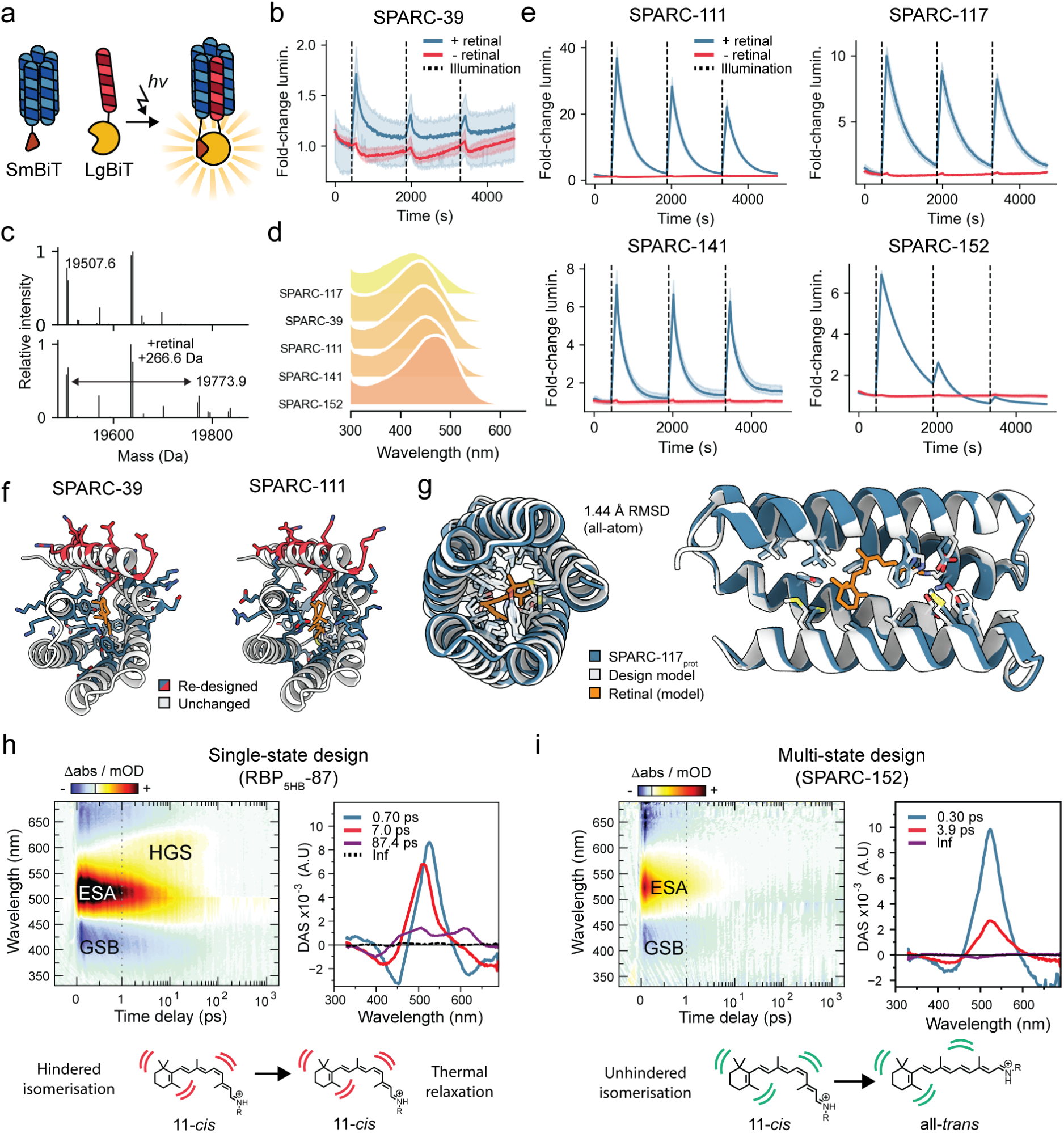
Experimental characterisation of *de novo* designed light-activated heterodimers. **a,** Schematic of the assay for screening light-activated heterodimerisation using a split-luciferase reporter. **b,** Split luciferase assay of the initial hit SPARC-39. Shaded area is standard deviation of n=3 independent measurements. Vertical dashed line indicates illumination at 480 nm. **c,** Native MS measurement of retinal binding to SPARC-39. **d,** Absorbance spectra of SPARC-39 and optimised derivatives. **e,** Split luciferase assay of the four optimised designs SPARC-111, 117, 141, and 152. Shaded area is standard deviation of n=3 independent measurements. Vertical dashed line indicates illumination at 480 nm. **f,** Model of SPARC-111 showing positions that were re-designed from SPARC-39 in blue/red, and consistent positions in white. **g,** Crystal structure of SPARC-117 (blue, dark state) superimposed onto the corresponding design model (grey, AF3 prediction). **h and i,** Ultrafast spectroscopy measurements of retinal activation in RBP_5HB_-87 **(h)** and SPARC-152 **(i)**. Transient absorption spectra (left) and decay associated spectra (right) for each protein showing formation of an excited state absorption (ESA) state after light activation, and corresponding ground state bleach (GSB).

Overall, these data indicate that SPARC-39 functions as a light-activated heterodimer as designed. However, considering that the observed response was modest, we set out to optimise the hit computationally. From models of the two conformations of SPARC-39, we repeated the multistate sequence design and filtering while allowing more positions in the switch to be designed, reasoning that sampling second and third shell residues may improve conformational switching. From this round of computational optimisation, we ordered 60 optimised designs for experimental testing, of which 23 designs showed an increase in luminescence on illumination similar to or greater than the parent design (Supplementary Fig. 9). Further, 4 designs showed a greater than 7-fold increase in luminescence on

illumination when tested as purified protein, with the strongest response from SPARC-111 giving a greater than 30-fold increase (Fig. 3e, Supplementary Fig. 10). Since we observed a drastic increase in light-activation between SPARC-39 and SPARC-111, we further analysed the mutations performed. Alignment of the models showed that SPARC-111 was virtually unchanged from SPARC-39 (0.80 Å Cα RMSD). However, at the sequence level, 36 of 153 positions (23.5%) were changed between SPARC-39_prot_ and SPARC-111_prot_, and 10 of 26 positions in the peptide (38.5%). Mapping to the model showed that the majority of the mutations are found in the retinal binding pocket or helices involved in peptide binding, but with some more distal mutations in the helical cap to the SPARC_prot_ (Fig. 3f).

To validate the designs, we again used MS to confirm accurate retinal attachment at the designed position, and mutation of the core lysine or assay without retinal completely attenuated light activation (Supplementary Fig. 11). Interestingly, the designed SPARCs had no reduction in thermostability compared to RBPs, with no change in the secondary structure of SPARC-111 up to 90 °C (Supplementary Fig. 12). Further, to validate the structural accuracy of the designs, we solved the crystal structure of SPARC-117_prot_ loaded with 11-*cis*-retinal (Fig. 3g, Supplementary Table 1). This confirmed the high structural fidelity of the design in the dark (inactive) conformation, with a Cα RMSD of 0.92 Å (1.44 Å all-atom) to the design model. Unfortunately, 11-*cis*-retinal was not clearly distinguishable in the electron density map, possibly being partially lost due to photodamage or Schiff base hydrolysis during crystallisation

We next asked whether the multistate design performed on the SPARC proteins had made any detectable changes to retinal isomerisation. To this end, we examined the top four switching designs and the parent non-switching design by ultrafast spectroscopy. The closest natural analogues to the designed proteins would be the visual rhodopsins, such as bovine rhodopsin, which also use 11-*cis*-retinal as a chromophore. It displays several photointermediates in the femto- to nanosecond timescale: photorhodopsin at ∼570 nm, bathorhodopsin at ∼540 nm and a blue shifted intermediate at ∼480 nm^38^.

Examination of the non-switching design instead revealed only a long-lived excited state absorption (ESA) and an accompanying ground state bleach (GSB) turning into a hot ground state (HGS; Fig. 3h). These spectral signatures are reminiscent of the previously designed soluble analogue of Bovine rhodopsin, which was also inactive for switching, and suggests hindered retinal isomerisation in the single-state design^29^. In turn, the four multi-state designs produced far shorter ESA lifetimes, and an infinity lifetime showing a depletion of the 480 nm signal with a rise around 400 nm matching retinal in solution, which together suggest successful retinal isomerisation. (Fig. 3i, Supplementary Fig. 13). Taken together, these data indicate that single-state designed RBPs bind retinal, but are trapped in a single conformation, effectively preventing isomerisation. Meanwhile, the subsequent multistate design campaign was successful in generating proteins that do not restrict retinal isomerisation and subsequent peptide binding.

### Rational engineering of designed light-responsive proteins

One of the limitations of existing light-responsive proteins is that they are challenging to modify beyond their existing properties or mechanisms^39^. With designed light-activated heterodimers in hand, we asked whether this system was amenable to customisation and tuning of their properties. To this end, we designed constructs to create orthogonal light-activated heterodimer pairs, shift the activation wavelength towards longer wavelengths, and to increase the sensitivity of the PPI.

Initially, we investigated whether we could create two orthogonal light-activated heterodimers for parallel control. Pairwise assay of the top four assayed designs confirmed that none of these pairs were orthogonal, likely due to their high sequence and structural similarity (Supplementary Figs. 14 & 15). Instead, inspection of the 3D model for SPARC-111 showed a possible route to an orthogonal pair. The interface has clear charge patterning, with complementary salt bridges from four glutamates on the peptide modelled to interact with four lysines on the bundle (Fig. 4a). In a classic piece of protein engineering, rationally swapping the charges on this interface created a new heterodimer, SPARC-111_ortho_, without a decrease in light response, but where the PPI is now orthogonal to SPARC-111 (Supplementary Fig. 16).

**Fig. 4:**
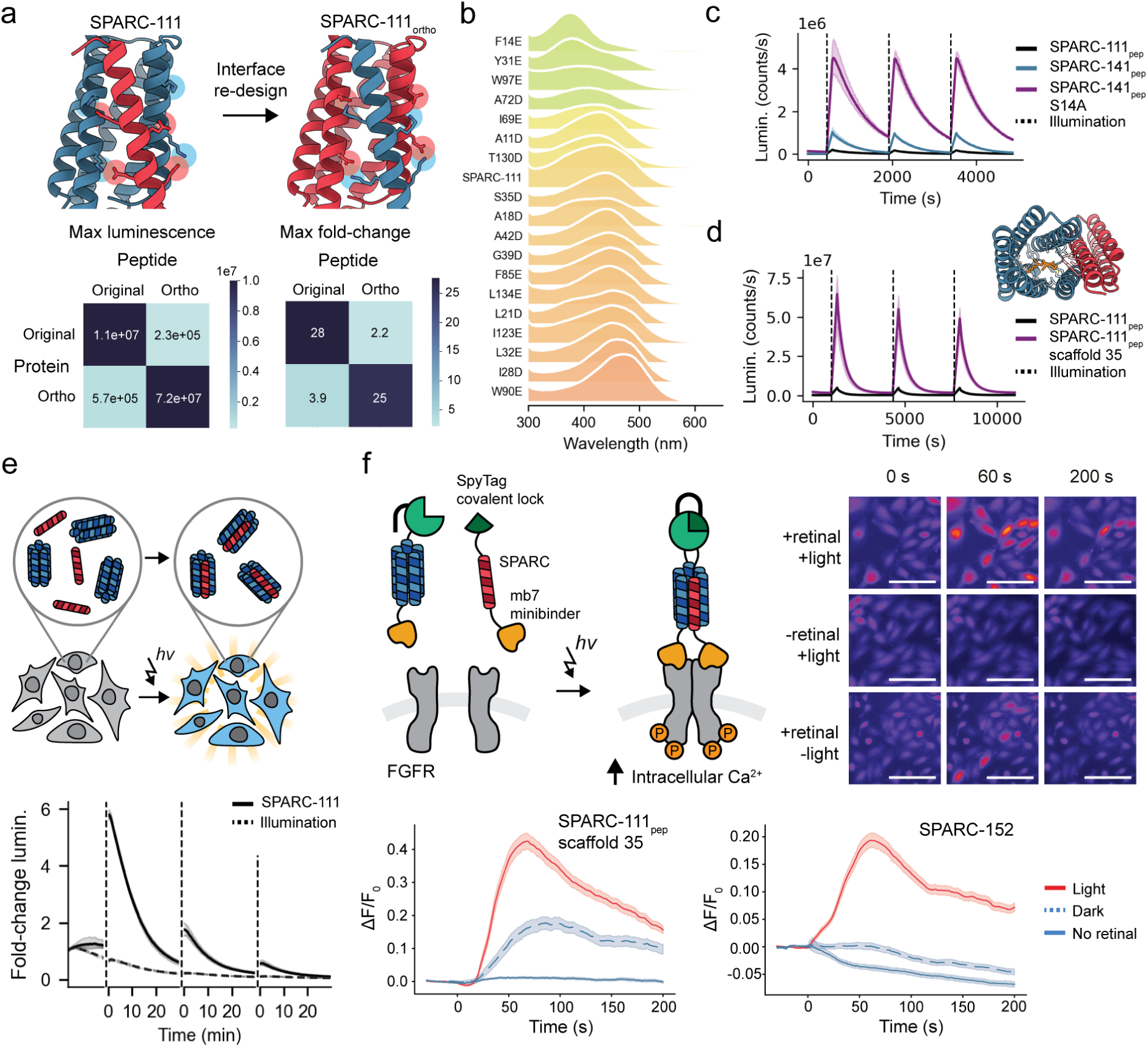
Rational improvements in light-responsive proteins and proof-of-concept applications in cells. **a,** Design models for SPARC-111 and SPARC-111_ortho_, and a split-luciferase orthogonality matrix. **b,** Absorbance spectra for SPARC-111 and point mutations in the retinal binding pocket. **c,** Split-luciferase assay of SPARC-111_prot_ combined with the original peptide (SPARC-111_pep_), SPARC-141_pep_, and the mutated SPARC-141_pep_-S14A. Shaded area is standard deviation of n=3 independent measurements. **d,** AF3 model for SPARC-111_pep_ scaffold 35, and split-luciferase assay of SPARC-111 with the original and scaffolded peptide. Shaded area is standard deviation of n=3 independent measurements. **e,** Split luciferase activation in HEK-293T cells. Shaded area is the standard deviation of n=3 independent biological replicates. Dotted line is the same constructs without added 11-*cis*-retinal. **f,** Engineering of light-dependent cell signaling agonists. Per-cell relative calcium levels are shown over time; SPARC-111_prot_ I28D + SPARC-111_pep_ scaffold 35 (top) and SPARC-152 (bottom). Protein is added at 0s. Mean of all cells per condition are shown as line and standard deviation is shown as shaded area. Per condition 3 to 4 biological replicates are shown. Representative microscopy images are shown to the right (SPARC-152), scale bars are 100 µm.

Next, we created mutations to SPARC-111_prot_ to shift the activation wavelength towards longer wavelengths. Addition of negatively charged residues around the Schiff base or β-ionine ring can both red-shift the retinal activation wavelength^40,41^. Screening point mutants of core residues in the bundle to glutamate or aspartate indicated two positions that produced a red-shifting in absorbance by up to 30 nm (Fig. 4b). Moreover, assay of the mutations for light activation indicated that one mutation, I28D, also improved the response of the switch, giving a 2-fold increase in luminescence with a similar fold-change (Supplementary Fig. 17).

Following this success, we set out to improve the sensitivity of the heterodimer. From the pairwise assay of the top four designs, we noted that the combination of SPARC-111_prot_ and SPARC-141_pep_ had a greater maximum luminescence upon illumination than the designed SPARC-111 (Supplementary Fig. 14). To investigate if we could improve the affinity of this interaction further, we made three mutations and their combinations to SPARC-141_pep_. One mutation, S14A, gave a 4.5-fold increase in max luminescence, which we attribute to the removal of an unsatisfied hydrogen bond in the interface and slight improvement of hydrophobic contacts (Fig. 4c, Supplementary Fig. 18). Moreover, when combined with the I28D mutation to SPARC-111_prot_, this combination has a 95-fold increase in luminescence compared to the original SPARC-111 pair, while retaining a similar or greater fold-change (Supplementary Fig. 19).

Finally, we asked whether the peptide component of the heterodimer could be scaffolded as part of a folded globular protein. We previously attempted this with SPARC-39, but the poor response of this switch meant that none of the scaffolded designs showed any light-response (Supplementary Fig. 9). Now, with the improved SPARC designs, we generated 40 miniproteins scaffolding SPARC-111_pep_ using RFDiffusion and ProteinMPNN_sol_ (Fig. 4d)^42,43^. Of 40 tested designs, more than half showed some degree of light-activated binding, with some showing greater signal and responsiveness than the parent design, indicating that scaffolding had improved the switching characteristics (Supplementary Figs. 20 & 21). Overall, these efforts confirm that the designed heterodimer system is amenable to modification with only a few hand-picked mutations, or by established protein design methods.

### Proof of concept for light-controlled processes in cells

We next asked whether the designed light-activated heterodimers could be used to control processes in living cells. First, we investigated whether the designed system could be used to activate PPIs in mammalian cells. To this end, we co-transfected the two components of the highest performing design, SPARC-111, fused to a split luciferase into HEK-293T cells (Fig. 4e). Illumination of the designed light-activated heterodimer in cells again resulted in an increase in split luciferase activity, confirming that the designed system can act as a light-responsive PPI in living cells (Supplementary Fig. 22). Moreover, this activation was retained over successive illuminations, though with a reduction in overall response which we attribute to a decay in the available luciferase substrate. Here however, incorporating the switch-improving mutations to the peptide or protein did not increase the luminescence signal in cells (Supplementary Fig. 22).

Next, we asked whether the SPARC system could be used to control complex cellular processes, such as cell signalling, in a light-dependent manner. To this end, we chose to investigate light-dependent activation of fibroblast growth factor receptor 1 (FGFR1). FGFRs are receptor tyrosine kinases (RTKs) with roles in cell growth, embryonic development, tissue repair, and certain cancers^44^. Dimerisation of FGFR leads to receptor autophosphorylation, Ras signalling, and ultimately intracellular calcium release (Fig. 4f)^45,46^. To activate FGFR1 in a light-dependent manner, we fused a previously designed minibinder (mb7) against FGFR1c to the two components of the SPARC system^47^. Thus, dimerisation of SPARC should induce FGFR1c dimerisation and intracellular calcium release, which we monitored with a calcium sensitive dye^48^. Activation with a constitutive dimer created by Fc fusion to mb7 confirmed that dimerisation was sufficient to initiate intracellular calcium release. However, initial investigations with the SPARC system showed little response, which we posited could be due to fast dissociation of the SPARC heterodimer leading to weak receptor activation. To overcome this limitation, we added the components of the SpyTag/SpyCatcher pair to the SPARC system^49^, in effect creating a covalent lock to prevent dissociation. Investigation *in vitro* determined that light activation was sufficient to increase the formation of the isopeptide bound complex (Supplementary Fig. 23). Addition of the SPARC-111-SpyTag system induced a weak calcium response only when retinal was present and following illumination. Moreover, testing of the improved SPARC combinations (SPARC-141_pep_-S14A and SPARC-111_pep_-scaffold-35) greatly improved the response (Supplementary Fig. 24). Thus, the designed light-activated heterodimers can be used to control both intracellular PPIs and cell surface receptor clustering in a light-dependent manner.

### Conformational dynamics of the SPARC switch

Having shown the fine-tuning of the SPARCs and their functionality in cellular systems, we sought to characterise the conformational dynamics underlying their switching mechanism. Central to the design hypothesis is a light-activated conformational change driven by retinal isomerisation from 11-*cis* to all-*trans*. Therefore, to investigate light-activated conformational changes in the designs, we examined them by nuclear magnetic resonance (NMR) spectroscopy with a custom fibre-optic cable coupled to an LED to illuminate the sample directly in the spectrometer and monitor the conformational changes occurring in solution in real time^50^.

We initially recorded 2D ^1^H,^15^N–heteronuclear single-quantum coherence (HSQC) NMR spectra for all of the top-performing SPARC_prot_ designs without retinal (SPARC-111, 117, 141, and 152; Supplementary Figs. 25, 28, and 29). All 4 designs yielded spectra with extensive chemical shift dispersion indicative of well-folded states, yet SPARC-111 and 152 displayed overall the sharpest crosspeaks and most homogeneous intensities, and were therefore subject to deeper investigation. Loading these SPARC_prot_ designs with 11-*cis*-retinal resulted in pronounced crosspeak shifting and sharpening, indicative of a structural response of the proteins to ligand binding and a reduction of conformational dynamics (Supplementary Figs. 25, 28, 29).

To measure light-activated conformational changes in these designs, we illuminated the 11-*cis*-retinal loaded samples *in situ* and collected a series of short (2.5 – 5 minutes) SOFAST-HMQC ^1^H,^15^N spectra in an attempt to capture structural changes taking place soon after light activation. Light activation resulted in extensive chemical shift perturbations relative to the pre-illumination spectra, indicative of a conformational change, but we did not capture intermediate conformations during the time series after illumination (Supplementary Figs. 26, 27). Next, we asked whether the observed perturbations could be attributed to retinal isomerisation from *cis* to *trans*. Measurements of the SPARC proteins loaded directly with all-*trans* retinal overlayed precisely with the light activated sample, confirming that the observed conformational response can be attributed to retinal isomerisation (Supplementary Figs. 28, 29). Moreover, in the dark state both SPARC proteins showed sharp, single-species crosspeaks of homogenous intensities, suggesting a stable structure with limited dynamics (Fig. 5a, b, Supplementary Table 2 and Figs. 28, and 29). In turn, after light activation the chemical shift perturbations are accompanied by broadening and splitting of numerous crosspeaks, indicative of increased conformational dynamics in the intermediate and slow chemical shift timescales (Fig. 5a, b, Supplementary Table 2 and Figs. 26, 27, 28, 29).

**Fig. 5:**
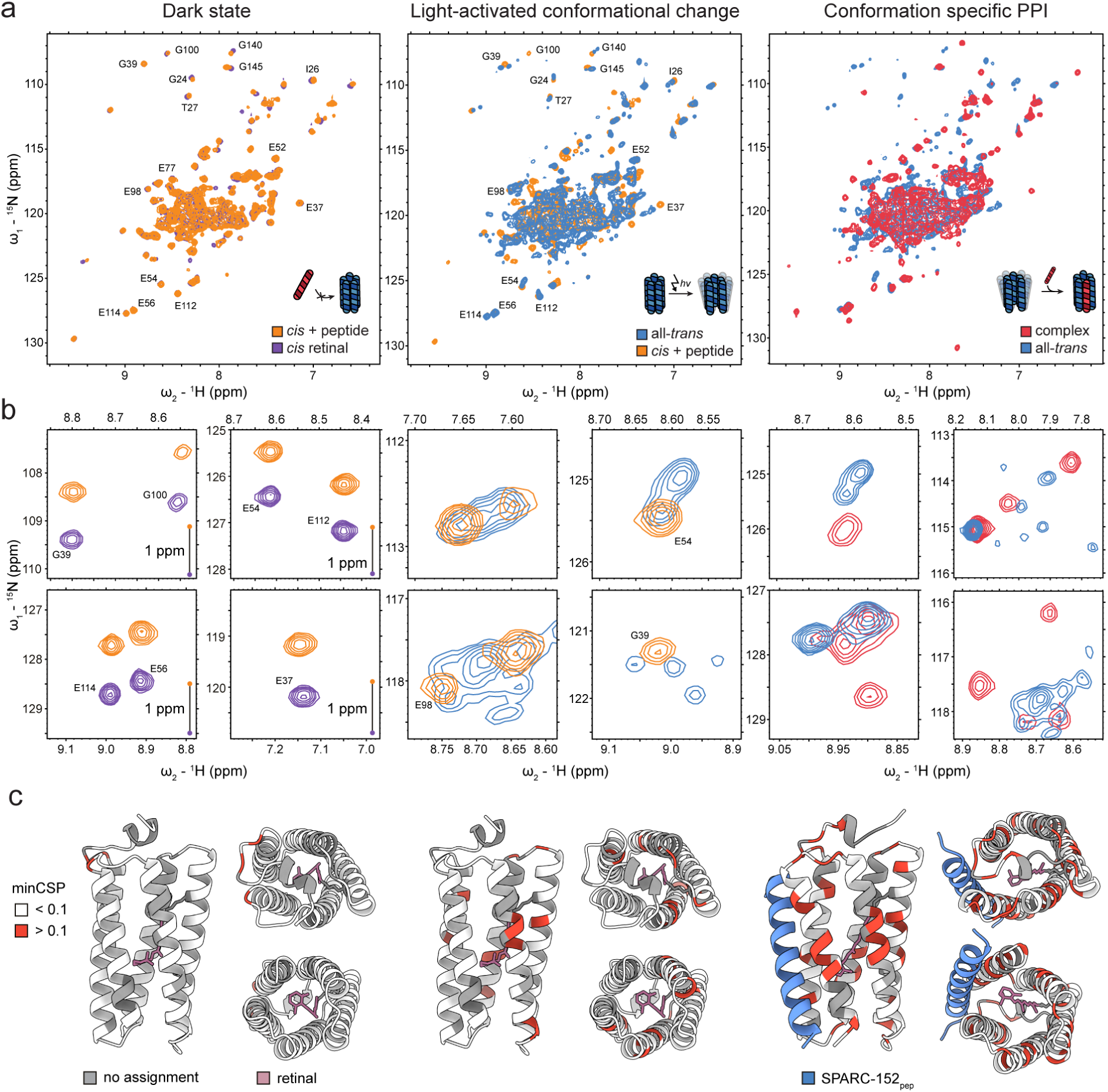
Following SPARC conformational dynamics with light-activated NMR. **a,** ^1^H,^15^N HSQC spectra of SPARC-152 following the steps of light-activated binding. Samples were prepared by loading SPARC-152_prot_ with 11-*cis* or all-*trans*-retinal and optional addition of SPARC-152_pep_. Corresponding *in situ* illumination experiments are presented in Supplementary Figs. 26 and 27. Isolated crosspeaks are annotated with their assigned residues. Assignments were obtained for SPARC-152 loaded with 11-*cis*-retinal (purple). Where possible, assignments were transferred to the corresponding sample with peptide (orange). **b,** Enlarged views of representative crosspeaks from **a**. In the dark state comparisons (left) the HSQC of 11-*cis*-retinal loaded SPARC-152 (purple) has been shifted by +1 ppm along the ^15^N dimension to allow for a better comparison of overlapping crosspeaks. **c,** Chemical shift perturbations mapped to structure models (minimal CSP, calculated relative to the 11-*cis*-retinal-loaded protein). From left to right: dark state (11-*cis*-retinal + peptide), light-activated (all-*trans*-retinal), and light-activated plus peptide. Residues without resonance assignments are coloured in grey, retinal is coloured in purple and the peptide is coloured in blue.

We then repeated the experiment but with added cognate peptide (SPARC-111_pep_ or SPARC-152_pep_). Measurement before illumination showed no significant changes to the spectra relative to the samples without peptide, confirming no interaction with the inactive, dark state structure (Fig. 5a, b). Yet, after light activation *in situ*, we observed widespread chemical shift perturbations distinct and larger than those displayed by the light-activated sample without peptide, indicative of light-activation accompanied by peptide binding (Supplementary Figs. 26, 27). Moreover, this sample showed an overall decrease in the number of crosspeaks and more homogenous intensities, indicating a partial suppression of dynamics in the complex (Fig. 5a, b). Again, we did not observe intermediate conformations upon light illumination but the observed spectra matched the spectra recorded on all-*trans* loaded SPARC incubated with the respective peptide (Supplementary Figs. 26, 27, 28, 29). Moreover, we confirmed that direct incubation with all-*trans*-retinal indeed increased the interaction with the peptide (Supplementary Fig. 30).

Thus, with *in situ* light activation we are able to induce and observe structural changes in the designed light-responsive proteins. To identify which areas of the protein undergo conformational changes in response to light, we assigned the backbone resonances for SPARC-152_prot_ loaded with 11-*cis*-retinal. Calculation of the minimal chemical shift perturbations to the same sample with added peptide again showed minimal changes. Next, we evaluated the structural changes relative to the light-activated spectrum and the complex. In the design models, the conformational change is localised to the *N*-terminal helix of the 5-helix bundle, with the peptide binding between helices 1 and 5. The minimal chemical shift perturbations from 11-*cis* to light-activated protein point at widespread changes in chemical environment for many residues upon light activation, consistent with conformational change triggered by retinal isomerisation which become even more extensive on complex formation (Fig. 5c).

Thus, taken together, these data support the designed mechanism of a light-activated conformational change driven by the retinal isomerisation, where the protein transitions from more dynamic states (apo and all-*trans*-retinal) to more conformationally defined states (11-*cis*-retinal, peptide-bound) providing a picture of the conformational landscape of the SPARC switching mechanism. Our data indicate that in total at least four distinct conformational states of the SPARC designs can be observed (apo protein, 11-*cis-*loaded, all-*trans*-loaded, and all-*trans*-loaded protein bound to its cognate peptide) and that the equilibrium between those states is directly modulated by the retinal isomer.

## Conclusions

The design and engineering of stimulus-responsive proteins is a longstanding problem in computational protein design and synthetic biology. Stimulus-responsive proteins enable a biological process to be activated or deactivated on command, enabling a level of temporal control unattainable, for instance, by popular cellular perturbation approaches such as genetic knockouts. Protein engineering has shown great promise in exploiting natural stimulus-responsive proteins, yet *de novo* design promises unmet flexibility and application-tailored design rather than re-purposing natural systems^51^. The optogenetics toolbox has come to rely on a relatively small set of photoreceptors from nature^3^. These modules are powerful, but their properties, such as their size, expression, activation wavelength, photoswitching kinetics and dynamic range, are often fundamentally limited.

Here, we present the design of new-to-nature photoreceptors. We develop the SPARC system entirely from the bottom-up, designing chromophore binding, a light-activated conformational change driven by retinal isomerisation, and a conformation specific PPI into a single protein. We show that these designed proteins function as light-responsive heterodimers, and we demonstrate the light-dependent control of molecular interactions *in vitro*, and in cells. Overall, these designed proteins have built-in features that fulfil all the necessary functions of natural photoreceptors within minimal, designable scaffolds. We have shown that the SPARC system is amenable to engineering and enables tuning of properties like affinity, activation wavelength, orthogonality, and even the structure of the effector domain. Some of these properties are unavailable to current photoresponsive domains, such as the flavin-based LOV, BLUF or Cry2 systems, where their parallel use is effectively prohibited by their restriction to blue light activation^20^. Instead, retinal can absorb across the visible spectrum into the near UV^24,41^, which, if combined with the PPI orthogonality demonstrated here, could spawn optogenetic pairs that can be used in parallel in cells. Thus, the designed SPARC proteins have the potential to meet, and in some cases exceed, the capability of natural photoreceptors.

Central to the design strategy for the SPARC system is a ligand-dependent light-activated conformational change, unveiling a conformation-specific PPI. The design of proteins to bind predefined small-molecule ligands, as well as those that present engineered dynamic behaviors or conformational changes remain challenging in many accounts, and previous works have designed either protein dynamics^52^, or PPIs^8^, that respond to light. Here, we have integrated all these biochemical capabilities into a single scaffold, bringing designed proteins closer to the multifunctional proteins observed in nature. Using NMR and light-activation *in situ*, we are able to track changes to the conformational landscape. Overall, we observe four distinct conformational states in the designed proteins, with appreciable changes in dynamics that match the model of a transition from a well-resolved dark state to a dynamic, conformationally flexible light-activated intermediate, that regains some conformational selection upon peptide binding. Moreover, we have shown that these dynamics are essential to the light-response, with single-state designs instead restricting retinal isomerisation due to structural inflexibility. Therefore, the SPARC system is a strong indicator that designing proteins that can access multiple conformations is not only tractable, but critical to unlocking new protein functions. Indeed, the design logic developed for SPARCs (coupling a stimulus-responsive small molecule to protein conformational changes) could be applied to enable protein designers to create new multistate proteins with prosthetic groups responding to a variety of environmental cues.

Part of the versatility of the SPARC design comes from the use of retinal as a light-responsive moiety. Retinal in membrane-bound rhodopsins is the foundation of optogenetics, with enormous impact in neuroscience^53^. Retinal is therefore biocompatible and accessible for *in vivo* applications^54^. SPARCs naturally incorporate retinal, circumventing the need to use light-responsive non-canonical amino acids via genetic code expansion^8^. Therefore, the SPARC system is well-aligned with the necessary requirements to develop from proof-of-concept into a genuine tool for optogenetics in broader contexts in biology.

Despite the successes described here, the SPARC system still has limitations to be overcome. The wavelength shifting mutations are still insufficient to achieve orthogonality of activation, and the majority of the tested proteins still absorb at shorter wavelengths than those found in rhodopsins. The efficacy of the SPARC system in cellular systems is also not yet on par with contemporary optogenetic tools, most notably requiring a covalent lock to aid activation of cell receptors, and further work will be required to explore their wider functionality in cells. Finally, it is perhaps unclear how available retinal is to the soluble proteome due to its solubility, but could be mitigated by use of hydroxy-retinal derivatives^55^.

Nevertheless, the SPARC system marks the establishment of a platform for the development of *de novo* light responsive proteins that can be customised for diverse applications. Future work could improve the characteristics of the system, creating red-shifted activation wavelengths for improved penetration, and improving the signal-to-noise of the switch. Moreover, we envision that the SPARC designs could be engineered into new mechanisms of light responsiveness, such as the development of light-activated homo-oligomers for optogenetic clustering, or minimal *de novo* allosteric domains. Ultimately, we envision that these designed photoreceptors can become part of a new toolbox to explore and control biology with light.

## Methods

### Computational methods

#### Computational design of retinal binding proteins (RBPs)

For the design of *de novo* RBPs, we sampled minimal, designable folds with internal cavities. Retinal is a large hydrophobic molecule that requires complete or partial exclusion from solvent for Schiff base formation and to enable its full spectral properties. Therefore, we selected antiparallel coiled-coil topologies with 4, 5 and 6 helices and a beta barrel as our starting points. To create initial folds that suited our purpose, we began from experimentally validated crystal structures for a 4-helix bundle (8A3K)^56^, 6-helix bundle (8QAE)^25^, and beta barrel (32TF). As an experimentally validated structure for a single-chain antiparallel 5-helix bundle was not available, an AF3 model for one was created by prediction of the sequence for 8QAE without the final helix. As we intended to pursue dynamics, we reduced the helical length for the 5- and 6-helix bundles from 4 to 3 heptads (28 to 21 residues) to reduce the size of the hydrophobic interface, while the 4-helix bundle was tested as both 3 and 4 heptad variants. Finally, as we aimed to create a solvent excluded pocket and the 5 and 6-helix bundles have a continuous pore, we created a short *C*-terminal helical cap on the end of both these folds by generating an additional 10 residues with RFDiffusion from gaussian noise^42^.

To each of these starting folds, we generated a library of 500 diversified backbones with RFDiffusion in partial diffusion mode (diffuser.partial_T=15). Rather than designing ligand binding directly onto these scaffolds, we have generally found AF models more designable than RFDiffusion backbones. Therefore, we generated 10 sequences for each backbone using SolubleMPNN^43^, and predicted them with AF2 in single sequence mode (no MSAs), selecting only the best sequence per backbone (highest pLDDT, lowest RMSD to the RFDiffusion structure) to continue with. Backbones with no sequences scoring pLDDT > 0.8 and RMSD < 1 Å were rejected.

To each backbone we used Rosetta MotifGraft to add the essential lysine and glutamate pair for retinal protonated Schiff base formation^57^. Residues E113 and K296 from the structure of bovine rhodopsin (1GZM)^58^ were grafted onto the scaffolds using a solvent accessible surface area (SASA) filter to select for grafts where the residues had been inserted into the core of the structure. Next, the remainder of the protein core was mutated to alanine and 11-*cis*-retinal covalently docked onto the grafted lysine using Gnina^59^, generating 10 poses. Poses were filtered by radius of gyration (ROG) and centre of mass measurements to remove poses where retinal had been docked partially outside of the protein. Following, the residues in the core that had been mutated to alanine were redesigned around the docked retinal using LigandMPNN^34^, generating 50 sequences per pose. Here, LigandMPNN appeared to frequently make unintuitive choices for amino acids, potentially due to the presence of charged Lys and Glu residues in the protein core. Thus, to prevent clearly misguided residue choices we excluded cysteine and charged residues (CDEHKR), with a slight additional bias towards large hydrophobics to encourage packing (W:2.5, Y:2.0). Finally, these sequences were predicted using both AF2 and AF3 in single-sequence mode, selecting designs with pLDDT of the ligand >70, and high ligand consistency to the docked pose.

#### Sequence novelty analysis

Natural rhodopsin sequences were collected by searching the UniRef100 database for sequences annotated as ‘rhodopsin’^60^. UniRef100 was accessed on 22.09.2026 and the rhodopsin cluster comprised a total 71,663 sequences. Sequences were divided into microbial and visual (animal) rhodopsins by taxonomy, where all sequences with taxonomy IDs falling under the Metazoa subtree were labelled as visual (animal) rhodopsins and all other sequences annotated as microbial rhodopsins. Sequences shorter than 50 residues or longer than 800 residues were excluded. In total this resulted in 60,273 sequences annotated as microbial rhodopsins, and 11,365 annotated as visual rhodopsins. In addition, the 27 experimentally validated *de novo* proteins (RBPs and SPARCs) were included.

Amino acid sequences for all of the natural and *de novo* proteins were then encoded using the ESM2 protein large language model, a 650 million parameter model trained on the UniRef50 database^31^. Sequences were passed through the model to generate per-residue representations, which were mean-pooled across all sequence positions to produce a fixed-length 1,280-dimensional embedding vector for each protein.

All of the computed embedding vectors were then projected to two dimensions using the Uniform Manifold Approximation and Projection (UMAP) algorithm with cosine distance as the metric. The 27 *de novo* proteins were embedded jointly with the natural sequences so that their positions in UMAP space reflect their true distances from the natural rhodopsin sequence landscape.

#### Computational design of light-responsive proteins (SPARCs)

To design light-responsive proteins coupled to retinal isomerisation, we started from AF3 models of experimentally validated RBPs. All of these proteins had been designed to bind 11-*cis*-retinal, and so their mechanism of activation would be a *cis* to *trans* isomerisation.Retinal was converted to the all-*trans* isomer in each RBP model using RDKit, aligning the new ligand on coordinates of retinal C13, C14, C15, and C20 atoms. Models in which this produced a clash with the backbone without disrupting the protein fold were taken forward as starting points for a conformational switch.. One design(RBP_5HB_-3) appeared particularly suited:its Schiff base linkage sits near the top of the 5-helix bundle so that isomerisation would be efficiently transferred to the protein in between the *N*- and the *C*-terminal helix of the bundle. As a result we chose to optimise this sequence using LigandMPNN to repack the core, generating a further 13 sequences from the same backbone.

From these starting points, retinal was isomerised using RDKit as described. Then RFDiffusionAllAtom was used to diffuse the part of the structure (helices 1 and 5) clashing with retinal to move the protein and resolve the clash^32^. Simultaneously, an additional 25-30 residues were diffused from gaussian noise to generate the new helical peptide binder. As the publicly available version of RFDiffusionAllAtom cannot accommodate multiple protein chains, these residues were diffused as an *N*-terminal extension to the protein and the loop deleted in post-processing. Designs were filtered at this stage to ensure the *N*-terminal extension was structured (assigned as helical with a continuous length of at least 22 residues by DSSP assignment), that there was a sufficiently large structural change in the protein without disruption of the fold (RMSD between 2 and 13 Å, LDDT between 0.6 and 0.7). In addition, we filtered out designs which still contained a clash with the ligand (clash score < 0.05), and selected for designs that had sufficient contacts between the original protein and the new binder helix to present a likely protein-protein interaction interface (more than 120 Cα contacts).

Models of the dark state (5-helix bundle with 11-*cis*-retinal) and active state (peptide-bound 5-helix bundle, all-*trans*-retinal) were then used for multistate sequence design with ProteinMPNN or LigandMPNN, tying positions between the two states of the 5-helical bundle such that a single sequence was designed for both conformations^11,33^. Only residues that changed environment between the dark and light conformations were redesigned, as measured by a local distance difference test (LDDT), and the remainder were fixed from the experimentally validated protein. The positions for the 5-helix bundle were tied equally between the active and inactive states, while the peptide was only present in the active structure. If LigandMPNN was used for sequence design the core of the protein was also designed to increase diversity. On average, this resulted in 31% of positions being designed with ProteinMPNN and 53% with LigandMPNN. Sequences were predicted with both AF2 and AF3 with and without the peptide binder, to create models for the active and inactive conformations. Designs were evaluated for standard metrics (pLDDT > 80, iPTM > 0.7), and additionally for changes to the retinal isomer state (*cis* to *trans*) and for a conformational change to the 5-helix bundle upon interaction (4 – 6 Å Cα RMSD between dark and light state).

#### Design of SPARCs with scaffolded helices

To scaffold SPARCpep into folded protein domains, RFDiffusion was used to generate 3- or 4-helical structures incorporating the peptide in its bound conformation, starting from the SPARC_prot_–SPARC_pep_ complex. The peptide backbone and key binding-site residues were kept fixed, while additional segments were generated at the *N*-terminus, *C*-terminus, or both. Scaffold sequences were then designed using SolubleMPNN, preserving these binding-site residues and leaving the SPARC_prot_ sequence unchanged. Designs were evaluated using AF2 both as a monomer and in complex with SPARC_prot_, and selected based on prediction confidence and structural agreement with the designed models (pLDDT > 0.95, RMSD < 1 Å).

### Experimental methods

#### Cell culture

HEK 293T cells were maintained in DMEM without phenol red (Thermo Fisher Scientific, cat. no. 31053028) supplemented with 10% (v/v) fetal bovine serum (FBS, Sigma Aldrich, cat. no. SO615), 1X GlutaMAX (ThermoFisher Scientific cat. no. 35050038), 100 U/ml penicillin and 100 μg/ml streptomycin (Thermo Fisher Scientific, cat. no. 15140122). CHO cells stably expressing human FGFR1 isoform IIIc (CHO-R1c, gift from David Baker and Sangwon Lee) were maintained in F12-K medium (Thermo Fisher Scientific cat. no. 21127022) supplemented with 10% (v/v) FBS, 1% Penicillin-Streptomycin, and 10 μg/ml Puromycin. Cells were cultured at 37 °C and 5% CO_2_ in a humidified incubator and maintained below 80% confluency.

#### Protein expression and purification

Synthetic DNA fragments encoding the designed proteins were ordered from Twist Bioscience, or amplified from sequence-verified plasmids by PCR and inserted into custom bacterial expression vectors using Golden Gate Assembly as described previously to produce constructs with a *C*-terminal 6x His tag.^61^ All vectors used a ccdB counter-selection marker to remove background template. Vectors were amplified and isolated from *E. coli* HB101 and sequence verified (Microsynth). Sequence verified expression vectors were transformed into *E. coli* T7 Express (NEB). 250 ml LB expression cultures were inoculated 1:100 from overnight cultures and grown to OD 0.6 – 0.8 (37 °C, 220 RPM). Expression was then induced by addition of 0.5 mM IPTG and recombinant proteins were expressed overnight (18 °C, 220 RPM).

After expression, cells were harvested by centrifugation and the pellets resuspended in 30 ml IMAC wash buffer (50 mM HEPES pH 7.5, 500 mM KCl, 15 mM Imidazole) supplemented with 0.5 mg/ml Lysozyme, 0.02 mg/ml DNase I, and 1 mM PMSF. The resuspended cells were lysed by sonication, and the insoluble components removed by centrifugation (20,000 × g, 20 minutes) followed by sterile filtration. The clarified lysate was applied to a 0.5 ml Ni-NTA superflow IMAC resin (Qiagen) hosted in gravity flow columns (Bio-Rad), and the resin was washed twice with 10 column volumes (CV) of IMAC wash buffer. Bound protein was then eluted in 1 CV of 50 mM HEPES pH 7.5, 250 mM KCl, 500 mM imidazole. The eluted protein was further purified by size-exclusion chromatography (SEC) using either an S75 10/300 or S200 Increase 10/300 column (Cytiva), chosen according to molecular weight, in 50 mM HEPES pH 7.5, 150 mM KCl. Protein-containing fractions corresponding to the expected molecular weight were pooled and concentrated in appropriate spin concentrators. The purity of final protein preparations was assessed by SDS-PAGE and stocks were flash-frozen in aliquots in liquid nitrogen and stored at -80 °C until use.

#### Preparation of retinal stock solutions

Retinal stocks (10 mM) were prepared in the dark by dissolving 11-*cis-* or all-*trans*-retinal in ethanol and concentration determined from the absorbance at 380 nm using the literature extinction coefficients (11-*cis*: 24,935 M^-1^cm^-1^, all-*trans*: 42,880 M^-1^cm^-1^)^62^. Stocks were stored at -80 °C and handled in the dark under dim red light.

#### Colourimetric screening for retinal binding

The designed RBPs were screened for retinal binding by measurement of the colour change (red-shifting) of retinal absorbance, from 380 nm in solution to ≥440 nm, upon formation of a protonated Schiff base. Initial designs were expressed and tested directly as purified proteins to gather additional information on protein expression and solubility, while later designs were first screened in cell lysate and putative hits were validated as purified proteins.

For small-scale lysate screening, the designed RBPs were transformed into T7 Express *E. coli* and expressed in 5 ml LB expression cultures overnight (18 °C, 220 RPM, 0.5 mM IPTG) in 24-well plates. The cells were collected by centrifugation (25 minutes, 3,000 × g) and resuspended in 250 µl PBS-T supplemented with 0.5 mg/ml Lysozyme, 0.02 mg/ml DNase I, 1 mM PMSF. The cells were incubated for 1 hour at 4 °C shaking at 600 RPM, before clarifying by centrifugation (25 minutes, 3,000 × g). 100 µl of the supernatant was transferred to a black clear-bottomed 96-well plate and incubated with 30 µM 11-*cis*-retinal in the dark at 4 °C for 1 hour. An absorbance spectrum was then measured using a Tecan Spark plate reader measuring from 240 nm to 700 nm in 1 nm increments. Hits were selected for purification as designs that had an absorbance maximum or visible shoulder at 440 nm or greater.

For measurement of retinal binding to purified protein, 40 µM of the purified protein in 150 mM KCl, 50 mM HEPES pH 7.5 was incubated directly with 30 µM of 11-*cis*-retinal in a black clear-bottomed 96-well plate in the dark at 4 °C for 1 hour and measured with a Tecan Spark plate reader using an absorbance spectrum as described.

For an image of the colour-change of retinal binding to selected RBPs, 1 ml of retinal-RBP complex was prepared at 60 µM as described and placed in a transparent cuvette.

#### Intact mass spectrometry in native-like conditions

Samples were diluted in 50 mM ammonium acetate or injected directly in MilliQ water with an injection volume of 2 µL. The separation was performed on LC instrument (Vanquish Horizon, Thermo Fisher Scientific) using the NativePac OBE-1 SEC column (2.1 × 50 mm, 3 µm, Thermo Fisher Scientific) with isocratic flow of 50 mM ammonium acetate with flow rate 100 µL/min and column oven heated at 30 °C. Eluting proteoforms were analyzed using Exploris 240 Orbitrap FT-MS instrument operated in the intact protein mode with High Pressure option activated. Native mass spectra were acquired as full scans in the range 1500-8000 m/z at a resolution set to 15,000, RF Lens at 200 %, normalized AGC target of 300 %, maximum injection time of 200 ms, SID of 50 eV, while 10 microscans were averaged. The mass spectra were deconvolved using Biopharma Finder 5.3 (Thermo Scientific) with Respect algorithm.

#### Top-down mass spectrometry in denaturing conditions

The RBPs were analyzed via top-down MS approach by liquid chromatography (LC) high-resolution MS/MS at a number of different higher-energy collisional dissociation (HCD) energies to fragment the protein and produce fragment ions in order to localize the modification sites. Samples were injected as they were in MilliQ water with injection volume of 2 uL. The separation was performed on LC instrument (Vanquish Horizon, Thermo Fisher Scientific) using Aquity UPLC Protein BEH C4 pre-column (2.1 x 5 mm, 1.7 µm, 300 A, Waters) with flow rate of 400 uL/min and oven temperature set to 45°C. The following gradient of solvent B was used: from 15 to 50 % within 2.5 min followed by column washing and re-equilibration steps. Solvent A was composed of MilliQ water with 0.1% formic acid, while solvent B consisted of acetonitrile with 0.1% formic acid. Eluting proteoforms were analyzed on an Orbitrap Exploris 240 FT-MS benchtop instrument (Thermo Fisher Scientific, Bremen, Germany) operated in the intact protein mode with Low Pressure option activated, in positive polarity and using targeted MS/MS approach with ion multiplexing. For this, six precursor ions corresponding to different charge states of the same proteoform carrying retinal modification were input into the targeted inclusion mass list. The isolation window was set to 1.2 m/z, AGC target set as standard, maximum IT set as Auto, with 240,000 resolution at 200 m/z and averaging 10 microscans. Top-down MS analysis was repeated using 6 different values (between 20 – 50 %) of normalized collision energy (NCE). Top-down MS data were processed using Peak-by-Peak software with MS deconvolution workflow (Spectroswiss, Lausanne, Switzerland).

The resulting high resolution MS/MS spectra were deconvolved and the obtained neutral masses of b- and y-fragment ions were pooled together. Detected fragments were matched against the theoretical fragments of the designs, resulting from *in-silico* fragmentation of the target protein sequence with addition of +266.2035 Da (monoisotopic mass of retinal in the Schiff base, C_20_H_26_). Authorized fragment matching error was 1 ppm. Fragmentation maps were visualized with ProSight Lite (Kelleher research group, Northwestern University).

#### Circular dichroism spectroscopy

CD spectra were measured using a Chirascan V100 spectrometer with a 1 mm path length cuvette. Proteins were measured at 5 µM in 100 mM KCl, 10 mM HEPES pH 7.5. Full spectra measured ellipticity between 200 nm and 260 nm in 1 nm intervals, with a scanning speed of 20 nm min^−1^ and a response time of 0.125 s. A reference spectrum recorded with the same parameters and buffer at 5 °C was subtracted from the measured ellipticity. Measurements of ellipticity with respect to temperature (melting and cooling spectra), were recorded by measuring ellipticity at 222 nm every 1 °C from 5 – 90 °C and the reverse. Pre-melt and post-melt full spectra were collected before heating, and after cooling respectively. Ellipticity (deg) values were converted to mean residue ellipticity (MRE) (deg·cm^2^·dmol^-1^·res^-1^) by normalization to the number of peptide bonds in the protein, and the path length using the equation:

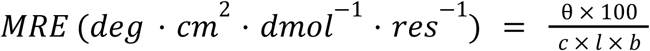

Where θ is the difference in absorbed circularly polarized light in millidegrees, c is the protein concentration in mM, l is the path length in cm, and b is the number of amide bonds in the protein.

#### Small scale split luciferase screening in lysate

For each light-activated heterodimer (5-helix bundle and peptide), the entire design was ordered as a single synthetic DNA fragment with two open reading frames separated by a ribosome binding site. Assembly of this fragment into custom expression vectors thereby created a bicistronic vector for co-expression of both components. Custom vectors were created such that insertion with Golden Gate assembly resulted in scarless fusion proteins of the bundle and peptide to the two halves of the split luciferase (LgBiT, SmBiT). Design models were used to decide on the orientation of the fusion constructs to maximise proximity of the fused split luciferase components upon complex formation, but the majority of the constructs were an *N*-terminal SmBiT fusion to the 5-helix bundle, and a *C*-terminal LgBiT fusion to the peptide.

To screen designs rapidly and cost-effectively, the gene fragments were assembled by golden gate assembly as described, and the reactions were transformed directly into T7 Express cells without sequence verification, using the ccdB counter-selection marker to remove background. The transformants were transferred into 1 ml LB cultures supplemented with 50 µg/ml Kanamycin in 2 ml 96-well deep well plates and grown overnight (37 °C, 220 RPM). The following day, 1.2 ml LB expression cultures in a fresh 96-well plate were inoculated with 50 µl of overnight culture and grown at 37 °C for 3.5 hours. Expression was then induced with 0.5 mM IPTG and the proteins expressed overnight (18 °C, 220 RPM).

The resulting cells were collected by centrifugation (25 minutes, 3000 × g) and either stored at -20 °C or lysed following one freeze-thaw cycle. For lysis, pellets were thawed and resuspended in 200 µl PBS pH 7.4 supplemented with 0.1% Tween-20, 0.5 mg/ml Lysozyme, 0.02 mg/ml DNaseI, and 1 mM PMSF. Cells were lysed for 1 hour at 4 °C while shaking at 800 rpm. The lysate was then clarified by centrifugation and 50 µL of supernatant was transferred into a 96-well reduced volume plate. 11-*cis*-retinal was added to a final concentration of 40 µM in a dark room under dim red light and lysates incubated for 60 minutes on ice. To account for high signal in the case of strong expression, each lysate was additionally diluted 1:5 and 1:25 in PBS. 10 µL of each sample was mixed with 10 µL NanoGlow luciferase assay substrate (Promega, prepared according to manufacturer‘s instructions) in white 384-well plates. Luminescence was then recorded on a Tecan Spark plate reader. The plate was first incubated for 10 minutes shaking (orbital pattern, 3 mm amplitude, 180 RPM) to reach a stable baseline. Following, luminescence was recorded for 1.5 minutes (50 ms integration time per well) to obtain a baseline in the dark. Wells were then illuminated with blue light using the fluorescence top reading mode (200 flashes, 485 nm ±20 nm excitation filter) to activate the retinal chromophore, followed by a luminescence measurement for 5 minutes (50 ms integration time per well) to record the activated luminescence. This was repeated three times for each well to record three successive illuminations of the same sample. To calculate a fold-change activation or normalised luminescence, the luminescence was divided by the mean luminescence from the final five recordings before illumination.

Plasmids encoding designs with a significant fold change in luminescence levels after illumination were transformed into *E. coli* HB101 and sequence verified. Assays were then repeated with sequence verified constructs for validation.

#### Split luciferase assay with purified protein

Purified SPARC_prot_ fused to SmBiT was first incubated with 11-*cis*-retinal in PBS pH 7.4 (30 µM SPARC_prot_, 40 µM 11-*cis*-retinal) for 1 hour on ice. Samples were prepared in a dark room under dim red light and protected from light as much as possible throughout. The retinal-loaded SPARC_prot_-SmBiT was further diluted and mixed with purified SPARC_pep_-LgBiT at equimolar concentrations (2, 1, and 0.5 µM). Protein solutions were prepared for luminescence measurement by mixing with Nano Glow luciferase assay substrate (Promega) in white 384-well plates. For each well, 10 µl of protein was mixed with 10 µl of substrate such that the final protein concentrations were 1 µM, 500 nM, and 250 nM in a 20 µl volume. Following, the purified protein was assayed in a similar fashion to the lysate screen in a Tecan Spark plate reader: after an initial shaking for 10 minutes (orbital pattern, 3 mm amplitude, 180 RPM) to reach a stable baseline, a starting luminescence was recorded in the dark for 7 minutes (50 ms integration time per well) followed by illumination with blue light using the fluorescence top reading mode (200 flashes, 485 nm (+/-20 nm) excitation filter) followed by luminescence recording for 20 minutes (50 ms integration time per well). The illumination and luminescence reading was repeated two times to record a total of three successive illuminations. To calculate a fold-change activation or normalised luminescence, the luminescence was divided by the mean luminescence from the final five recordings before the first illumination.

For optimised designs (e.g. SPARC-141_pep_-S14A) the signal at 250 nM was too high for accurate measurement and there was evident substrate depletion, so protein solutions were prepared at 100, 50, and 10 nM instead. For each assay the concentration is given in the figure legend.

#### Nano-Glo Live Cell Assay

HEK 293T cells were seeded in black 96-well glass-bottom plates (Corning, cat. no. CLS3904) at a density of 12,500 cells per well in 100 µL medium. After 24 hours, cells were transfected with 100 ng total DNA per well using 0.5 µL Lipofectamine 2000 (Thermo Fisher Scientific, cat. no. 11668019) according to the manufacturer’s protocol. Specifically, 25 ng of a plasmid encoding the respective SmBiT-SPARC_prot_ variant, 25 ng of a plasmid encoding the respective SPARC_pep_-LgBiT(D102Y) variant and 50 ng of pBluescript stuffer plasmid were co-transfected per well. As a positive control, 25 ng of a plasmid encoding a LgBiT(D102Y)-SmBiT fusion was co-transfected with 75 ng of the stuffer plasmid. The LgBiT(D102Y) mutant was chosen for its reduced activity upon NanoLuc reconstitution compared to the wild-type enzyme, leading to a lower rate of substrate depletion over the course of the assay.

The NanoGlo Live Cell Assay was performed at 22 hours post-transfection. All steps involving the Nano-Glo Live Cell Reagent and retinal were performed under dim ambient light in a darkened cell culture hood. Plates were protected from ambient light as best as possible during manual sample handling. First, the medium was aspirated and replaced with 100 µL Opti-MEM (Thermo Fisher Scientific, cat. no. 31985070) mixed with 25 µl Nano-Glo Live Cell Reagent (Promega, cat. no. N2012), prepared according to the manufacturer’s protocol. Next, 1.25 µL 11-*cis*-Retinal (1 mM in ethanol) was added per well to a final concentration of 10 µM. Controls were treated with 1.25 µL ethanol per well instead. After retinal addition, luminescence was measured in a Tecan Infinite MPLEX plate reader using the i-control software (version 2.0) with an integration time of 0.5 s at 37 °C. For all luminescence measurements, the transparent bottom of the plate was covered with black sealing tape (Thermo Fisher Scientific, cat. no. 236703) to prevent signal spillover into adjacent wells, which was intermittently removed for blue light illumination.

After 15 min of luminescence measurement in the dark, the cells were illuminated from below through the transparent well-plate bottom in a mammalian cell culture incubator using a custom setup with high-power blue LEDs controlled by a Raspberry Pi. Samples were illuminated for 1 min with a duty cycle of 5 s of light (30 μmol/m2/s; measured with a LI-COR light meter) followed by 10 s in the dark. Following illumination, luminescence was measured continuously for 30 minutes. Immediately after, samples were taken out of the plate reader for the next illumination phase in the incubator, followed by another 30 minutes of luminescence measurement in the plate reader as before. This cycle was repeated to a total of 3 light pulses and subsequent luminescence measurement. Since the exact time for sample transfer between the light pulse and luminescence measurement varied by a few seconds across the three biological replicates (manual sample transport step), the measurement time axes were aligned before averaging the three independent biological replicates. Each biological replicate represents the mean of three technical replicates, i.e. different wells transfected and treated in parallel.

#### Ultrafast transient absorption (TA) spectroscopy

Ultrafast TA spectroscopy experiments were performed using a home-built pump-probe setup described in detail elsewhere^63^. A Ti:Sa chirped pulse regenerative amplifier (Mai Tai SP-NSI, Spectra-Physics, Milpitas, CA USA) pumped by a Nd:YLF laser (Empower 45, Spectra-Physics, Milpitas, CA, USA) served as the fs-laser source for the setup. It was operated at a central wavelength of 800 nm and a repetition rate of 1 kHz, resulting in laser pulses with a pulse width of 100 fs. Excitation pulses were generated from a home-built two-stage noncollinear optical parametric amplifier (NOPA) with an interlayed prism compressor. For probing absorbance changes, the laser fundamental was focused into a 5 mm CaF_2_ crystal to generate supercontinuum pulses. The signal was detected using a custom spectrometer containing a sensor (MMS UV-VIS II, Carl Zeiss, Oberkochen, Germany) and preamplifying electronics (Tec5, Steinbach, Germany). The pump wavelength was adjusted to the respective absorption maximum, and pump and probe pulses were set to the magic angle (54.7°) configuration to eliminate anisotropic effects. Additionally, the sample was constantly moved in a plane perpendicular to the excitation beam to avoid multifold excitation and sample degradation. Samples were prepared in a 50 mM HEPES, 150 mM KCl buffer at pH 7.5 in a 1 mm cuvette. Absorption spectra were taken before and after each experiment to verify sample quality.

#### **X-** ray crystallography

Proteins were purified by SEC in 10 mM HEPES, 50 mM KCl, and incubated with 11-*cis*-retinal. Due to the limited solubility of retinal in aqueous buffers, the apo-protein was first incubated at 20 µM with 40 µM 11-*cis*-retinal in 50 mM KCl, 10 mM HEPES pH 7.5 before concentration of the holo-protein to crystallography concentrations.

Rod shaped protein crystals were grown for four weeks in the dark and harvested under dim-red light from a condition containing 0.2 M Lithium Sulfate 0.1 M Sodium acetate pH 4.5 and 50% PEG400 and flash-frozen in liquid nitrogen. Diffraction data were collected at Diamond Light Source beamline i24 (proposal mx42085) with a wavelength of 0.620 Å at temperature of 100 K. Data were processed on site using autoPROC. PhaserMR and Refmac were used for molecular replacement and structure refinement. A ligand-free model of the designed structure was truncated using the Chainsaw algorithm and used as template for molecular replacement in PhaserMR. The structure was refined in refmacat.

#### Nuclear Magnetic Resonance spectroscopy

All NMR experiments were performed on a Bruker spectrometer operating at a 800.13 MHz ^1^H frequency and equipped with a 5 mm CPTCI ^1^H/^13^C/^15^N cryoprobe controlled by an Avance Neo console. All samples were prepared in 20 mM MES pH 6.5, at 100 – 500 µM concentrations in 500 µL volumes to which 40 µL of D_2_O was added. All experiments were conducted at 308 K. All NMR spectra were acquired and processed using Bruker TopSpin 4.5. Isotopically labelled proteins were prepared by expression in M9 minimal media supplemented with ^15^NH_4_Cl and/or ^13^C glucose for ^15^N and ^13^C labelling respectively, and purified as described above.

^1^H,^15^N correlation spectra were acquired with either HSQCETF3GPSI (D1=1s) or SFHMQCF3GPPH^64^ (D1=0.2s) pulse programs, as indicated, taken off the Bruker Topspin library, and analyzed using Sparky-NMRFAM. For experiments illuminating the sample, we loaded the sample (approx 400 µL) into a 5 mm NMR tube with a built-in capillary that allowed insertion of the sanded end of the fibre-optic cable so that it could homogeneously illuminate the sample without actually touching the solution (this raised the level of solution to the equivalent of around 600 µl). To the fiber was coupled a 470 nm LED (14 mW, 1 A; Thorlabs M470F4) driven by a T-Cube LED Driver (Thorlabs LEDD1B). The fibre, capillary, and sample were inserted into the measurement chamber of the spectrometer prior to instrument setup (locking, matching, shimming) and measurement of the dark spectra. After collecting the dark spectrum, the sample was illuminated for 30 s in-place and new spectral acquisition was started immediately after shutting off the LED. Successive illuminations were attempted but did not result in further detectable changes to the spectra.

Experiments for backbone resonance assignment consisted of standard triple resonance spectra HNCA, HN(CO)CA, HNCO, HN(CA)CO, CBCA(CO)NH and HNCACB acquired on a 500 µM ^13^C,^15^N-labelled sample loaded with 11-*cis*-retinal^65^. Backbone assignments were obtained automatically with NMRtist^66^, a webserver implementation of the ARTINA pipeline^67^, and checked through manual inspection in the program CARA^68^. We assigned 80% of the backbone H,N correlations expected in the HSQC spectrum (85% excluding the 6x His-tag) and 85% of the CA and CB nuclei. Most of the missing assignments map to the surroundings of Lys76 (retinal binding residue), mainly in helix 4 and the *C*-terminal element that caps the helical bundle and in regions of helices 1 and 2 that are expected to be within reach of the retinal moiety according to our structural models.

Minimal chemical shift perturbations (mCSP) were computed relative to protein loaded with *cis*-retinal, for which assignments were obtained. Briefly, for each crosspeak assigned in the *cis*-retinal protein, we computed the distances to all crosspeaks in the target spectrum in CSP space:

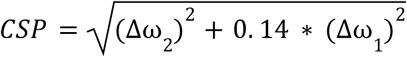

The residue was assigned with the CSP to the closest crosspeak in the target spectrum as mCSP.

#### Light-dependent FGFR activation

For light-dependent activation of FGFR, the SPARC system was combined with a minibinder (mb7) PDB ID 7N1J previously designed against FGFR1c (isoform IIIc). Chinese hamster ovary cells stably expressing FGFR1c (CHO-R1c) were seeded in 96-well plates and grown to 80-90% confluency. Cells were washed with DPBS (Thermo Fisher Scientific, cat. no. 14190144) and serum-starved for 3 h in F12-K medium. Cells were washed with DPBS and incubated with 5 µM Calbryte 520 AM (AAT Bioquest cat. no. 20651, prepared as 2.5 mM stock solution in anhydrous DMSO) and 2.5 mM Probenecid (AAT Bioquest cat. no. 20062) in Hank’s balanced salt solution (HBSS) for 30 min at 37 °C. Cells were washed with DPBS and 50 µL HBSS was added to each well. SPARC_prot_ variants were diluted to 20 µM in PBS and loaded with 40 µM 11-*cis*-retinal in the dark for 60 min on ice. SPARC_prot_ and SPARC_pep_ variants were then prepared at equimolar ratios and desired concentration in HBSS. Variants linked to the SpyTag/SpyCatcher system were mixed at 2 µM in HBSS and incubated for 10 min. either in the dark or under continuous illumination with a blue LED light. Protein solutions were added to the cells without further mixing, giving a final concentration of 1 µM of each component in the well. Calbryte 520 fluorescence was monitored using the GFP channel in widefield mode on a Nikon Crest X-Light V3 microscope with 20× objective, in a chamber maintained at 37 °C and 5% CO_2_, with an exposure time of 50 ms and a frame interval of 1 sec.

Baseline fluorescence was recorded for 30 s before protein addition, and imaging continued for 5 min after addition. For each recording, 300-400 cells were monitored on average.

The image series were analysed using BIOP-Desktop (v0.2.4, https://hub.docker.com/r/biop/biop-desktop?tag=0.2.4), using a Fiji-based pipeline combining pre-processing and segmentation steps. Time-lapse stacks were temporally projected (justified by minimal cell motility over the acquisition period) and square-root transformed (to enhance the detection of low-intensity cells). The resulting projected image was segmented using Cellpose-SAM to identify individual cells, generating a corresponding label image. This label image was then converted into regions of interest (ROIs), which were applied to the original time series to quantify cell-associated intensity at each time point (analysis script available on Zenodo: 10.5281/zenodo.23063068)^69^. The fluorescence was normalised per cell to the baseline recording and plotted over time using Python.

## Data availability

The crystal structure of SPARC-117_prot_ has been deposited in the protein data bank (PDB) under accession code 34XO. Design metrics and models of the proteins in this study, and backbone resonance assignments for SPARC-152 are accessible on Zenodo (10.5281/zenodo.23063068)^69^.

## Code availability

Code for the design of SPARC proteins is available on GitHub at https://github.com/LPDI-EPFL/sparc.

## Acknowledgements

We thank the National Eye Institute (NEI) for providing 11-*cis*-retinal. We thank F. Pojer, K. Lau and A. Larabi (Protein Production and Structure Characterization Core Facility, EPFL, Switzerland) for help with crystal screens and data collection. A.T.H. is supported by a Postdoctoral Fellowship from the Peter und Traudl Engelhorn Stiftung. R.R. is supported by an EMBO Postdoctoral Fellowship (ALTF 839-2026). D.N. was supported by the European Union (ERC, DaVinci-Switches, project number 101041570). B.E.C. is supported by the Swiss National Foundation, European Union. Views and opinions expressed are however those of the author(s) only and do not necessarily reflect those of the European Union or the European Research Council Executive Agency. Neither the European Union nor the granting authority can be held responsible for them. D.N. is also grateful for funding by the Deutsche Forschungsgemeinschaft (DFG, German Research Foundation) under Germanýs Excellence Strategy—EXC-3018/1—533587280. We thank the Swiss National Science Foundation (SNSF) for funding and support. We thank F. Praetorius (ISTA, Austria) for helpful discussions on designing conformational changes de novo. We thank A. Phal (Institute for Protein Design, USA), T. Schlichthaerle (Technical University of Munich, Germany), and M. Plum (EPFL, Switzerland) for helpful discussions about microscopy and monitoring intracellular calcium release. We thank J. Cross (University of Bristol, UK) for help with peptide synthesis. We thank A. Vinit, R. Guiet, and A. Seitz (Bioimaging and Optics Platform, EPFL, Switzerland) for help with microscopy setup and data analysis. We thank Dick Bradsell for continuous indirect support. We thank Aurélien Bornet and Anto Barisic at EPFL’s NMR facility for assisting in setting up the system for sample illumination inside the NMR magnet. X-ray diffraction data were collected at Diamond light source (DLS) beamline i24 (Proposal MX42085). We thank DLS for allocating beamtime and resources and the beamline scientists for technical support during data collection.

## Author contributions

A.T.H., N.M.G., and B.E.C. conceived the project and designed the experiments. A.T.H., N.M.G., C.L., and Y.K. developed the computational design methods and designed the proteins. A.T.H., N.M.G., C.L., K.P., E.O., S.G., and J.S. purified proteins. A.T.H., N.M.G., and C.L. performed the absorbance assay and split-luciferase experiments. C.L. and N.M.G. performed the FGFR activation experiments. L.A.A., N.M.G., and A.T.H. designed and performed the NMR experiments. N.A. and L.M. performed the mass spectrometry experiments. A.S.K., F.C.R., S.A, and D.N. designed and performed the in-cell split-luciferase experiments. R.R. and J.St. designed and performed the X-ray crystallography experiments. S.W. and J.W. designed and performed the ultrafast spectroscopy experiments. All authors contributed to data analysis and preparation of the manuscript.

## Competing interests

A.T.H., N.M.G., and B.E.C. are co-inventors on a provisional patent application 26 210 991.1 related to the proteins made in this study. The remaining authors declare no competing interests.

