## Supplementary Information for "De novo design of light-controlled conformational protein switches"

‡ Equal contribution.

### Contents:

#### Supplementary Figures 1 – 30

#### Supplementary Tables 1 & 2

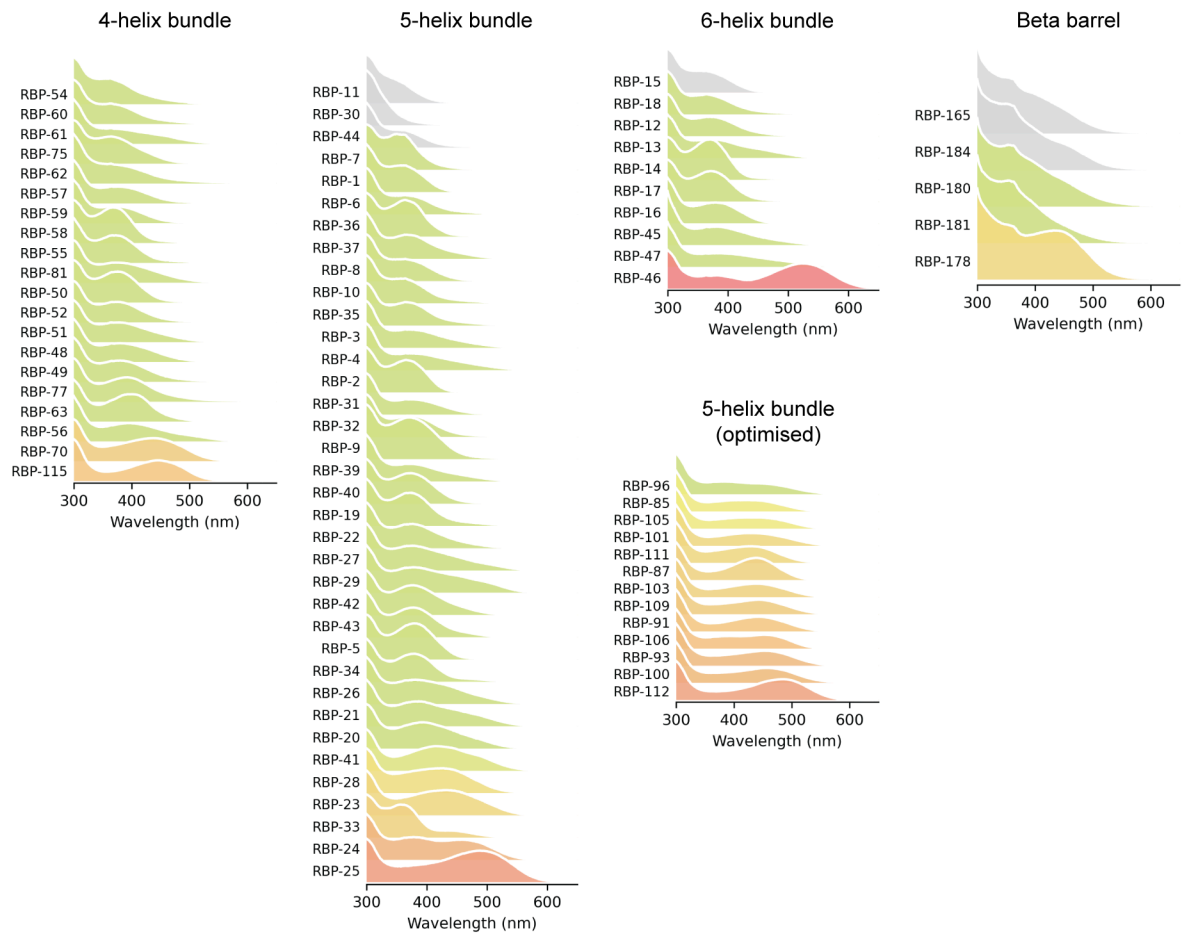

**Supplementary Fig. 1: Absorbance spectra of purified retinal binding proteins (RBPs).** RBPs incubated with 11-*cis*-retinal, coloured and ordered by their absorbance maximum.

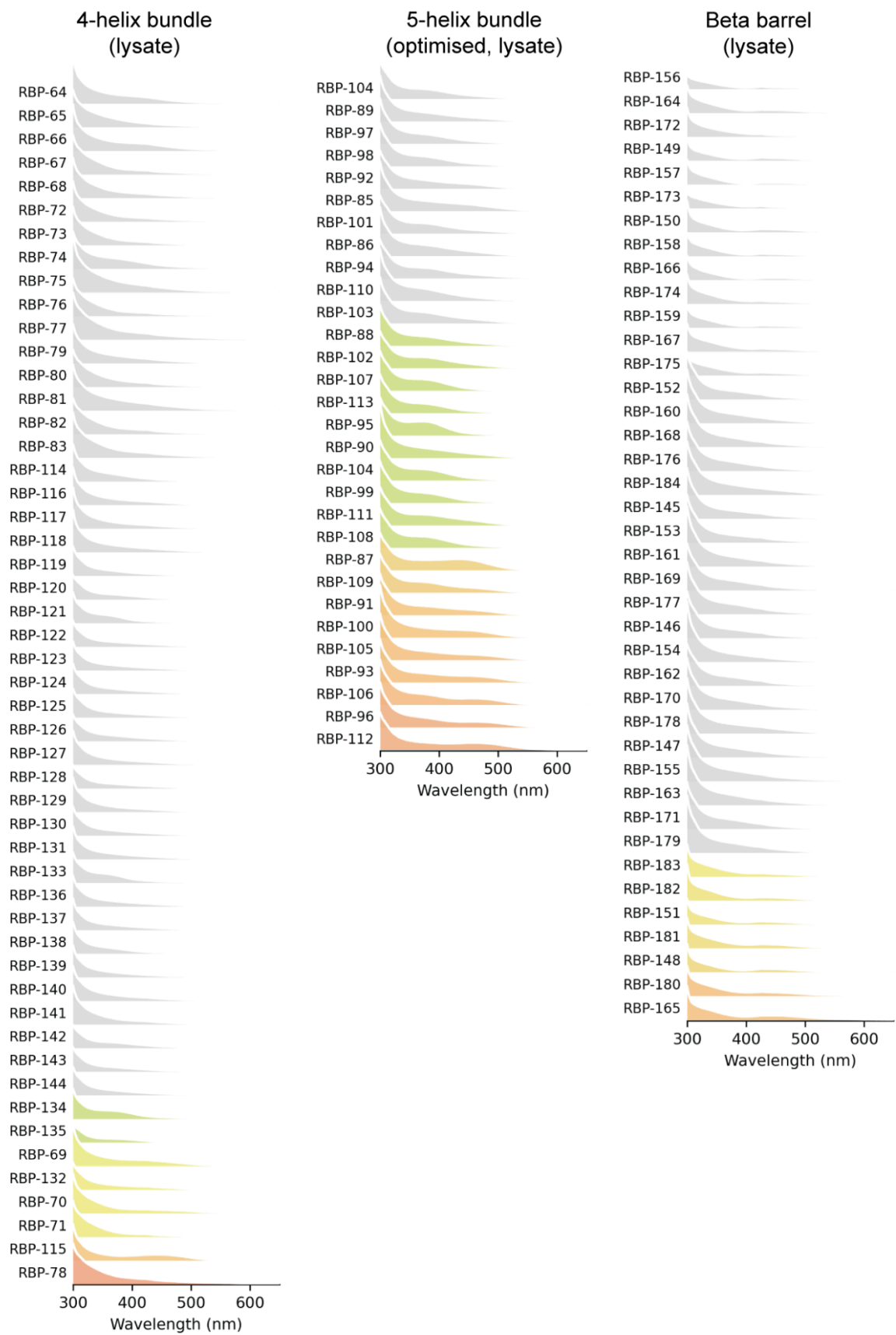

**Supplementary Fig. 2: Absorbance spectra of RBPs in *E. coli* lysate.** RBPs incubated with 11-*cis*-retinal, coloured by their absorbance maximum.

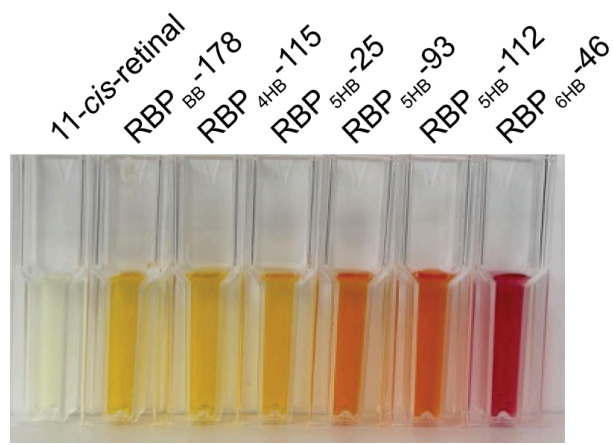

**Supplementary Fig. 3: Observed colour of purified RBPs.** RBPs incubated with 11-*cis*-retinal compared to free 11-*cis*-retinal in solution, all at 60 μM.

|  |  |  |  |  |  |
| --- | --- | --- | --- | --- | --- |
| RBP<sub>4HB</sub>-115<br>4-helix bundle<br>N S G K E L I E E A I K L A K E S T E S E E I V K E 25<br>26 L E K Y K K E S N S D K E K L D E I I K S L K E K 50<br>51 L E T G K E A I K F A E E N K D T L T K E S I K K 75<br>76 Y L L E E G E K S A E T T K R I V E T L K E I I E K K 100<br>101 S K D K E K I K E I I I E S L E R S L E T T K E L L I 125<br>126 E Y T K E L L L E G S H H H H H H C |  |  | RBP<sub>5HB</sub>-25<br>5-helix bundle<br>N S G K K W E E V I E L M K E E L E Y Y E A L A K G 25<br>26 E I D P V E A L E K I A E A K E K I L K L A V E A 50<br>51 L R E A G Y E E Y A A E L L E L A T A V I E A L R 75<br>76 K L A E A L K A G K P A V E Y L K E L I E Y Y R R 100<br>101 I V E A L R R L A E K A E D P E L L K E A A E E A A 125<br>126 E I I V E R I A K I F E D I L L E G K T E E A K K L K 150<br>151 E M M E G S H H H H H H C |  |  |
| <u>Precursor Mass</u><br>Type: Monoisotopic<br>Observed: 16,750.90<br>Theoretical: 16,750.92<br>Mass difference (Da): -0.017<br>Mass difference (ppm): -1.01 | <u>Scores</u><br>PCS: 965.70<br>P-Score: 2.7e-82<br>% Residue: 71% | <u>Modification</u><br>K50: 266.2 Da | <u>Precursor Mass</u><br>Type: Monoisotopic<br>Observed: 18,820.94<br>Theoretical: 18,820.95<br>Mass difference (Da): -0.002<br>Mass difference (ppm): -0.09 | <u>Scores</u><br>PCS: 1,564.72<br>P-Score: 8.3e-128<br>% Residue: 69% | <u>Modification</u><br>K40: 266.2 Da |
| RBP<sub>6HB</sub>-46<br>6-helix bundle<br>N S G K E L I E R L K E A I E R A I A L A R R V L E 25<br>26 E D L D E A E R L A L T A E A Y R A M A E A E R L 50<br>51 L A E L T K W P K I K E L L L R K A E L L E E M A 75<br>76 D L Y E R Y L A G E I S R E E Y L R A A A E L Y R 100<br>101 E L A E L L R A L A K L T K D E K L K E I L E R M 125<br>126 A E A A E E V A D L I D A L L A G E D V E A R L A 150<br>151 E A T R R A A E L G I E Y Y E Y W L L E L L K D E E 175<br>176 A Y E K Y L L E E Y A G S H H H H H H C |  |  | RBP<sub>BB</sub>-178<br>Beta barrel<br>N S G S N V V K V K F K L K V T D K E T G E V K E I 25<br>26 E G E A L V E K K G N V V K V D G E Y K V P G T D 50<br>51 E I R H F K A E G E L L S E E N G V T K Y H Y K L 75<br>76 L S E P P K E I E K E G D Y T L T E K E D K I E V 100<br>101 K G K D V D K E T G E E K E F T G E V T Y G S H H 125<br>126 H H H H C |  |  |
| <u>Precursor Mass</u><br>Type: Monoisotopic<br>Observed: 22,688.90<br>Theoretical: 22,689.89<br>Mass difference (Da): -0.994<br>Mass difference (ppm): -43.79 | <u>Scores</u><br>PCS: 1,690.89<br>P-Score: 2.4e-137<br>% Residue: 57% | <u>Modification</u><br>K67: 266.2 Da | <u>Precursor Mass</u><br>Type: Monoisotopic<br>Observed: 14,877.64<br>Theoretical: 14,877.63<br>Mass difference (Da): 0.017<br>Mass difference (ppm): 1.13 | <u>Scores</u><br>PCS: 1,588.53<br>P-Score: 1.3e-129<br>% Residue: 78% | <u>Modification</u><br>K85: 266.2 Da |

**Supplementary Fig. 4: Top down MS analysis of RBPs.** Fragmentation maps of one RBP from each designed fold. A blue line in each sequence marks a detected fragment from top-down MS, with a tail towards the *N*- or *C*-terminus indicating *b*- or *y*-fragments, respectively. Location of the 11-*cis*-retinal adduct (+266.2 Da) is indicated by an orange box. % Residue indicates achieved proteoform sequence coverage.

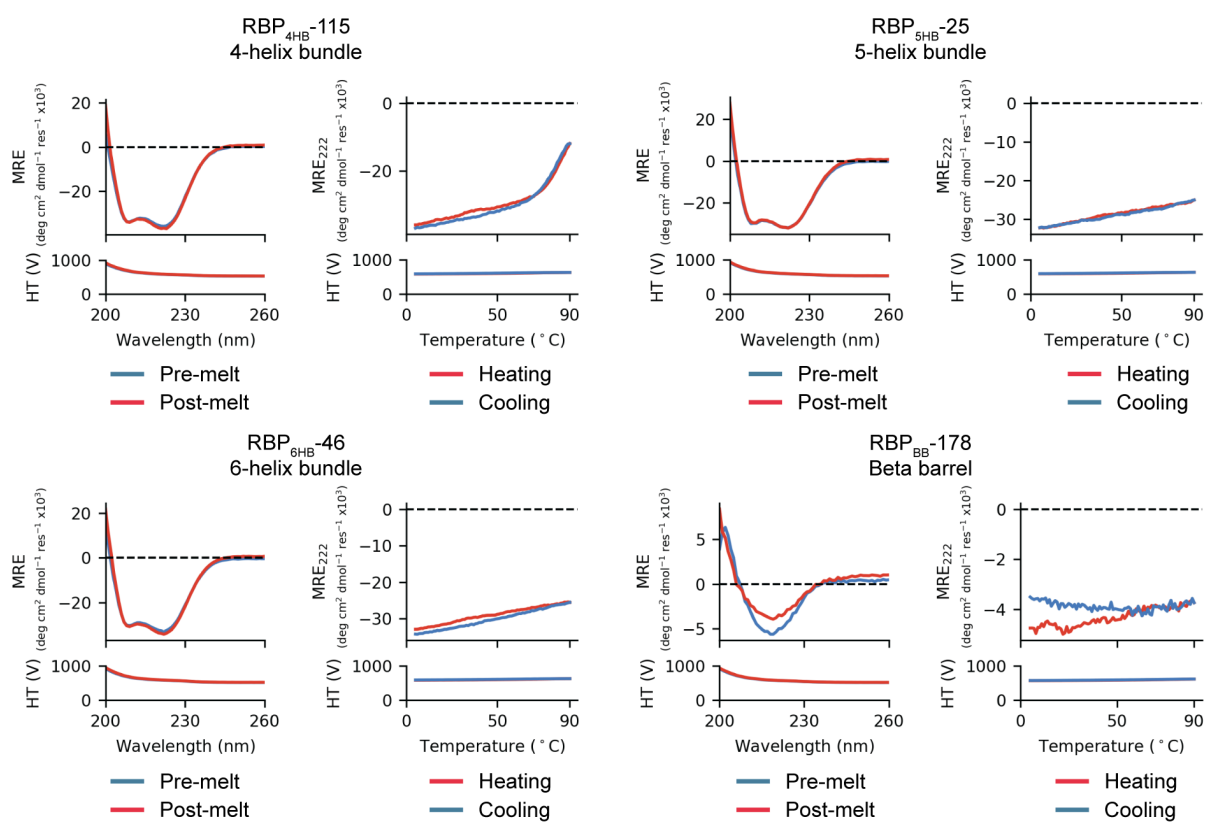

**Supplementary Fig. 5: Secondary structure and thermal stability of designed RBPs.** Measurement of secondary structure and thermal stability for each designed fold measured by circular dichroism (CD) spectroscopy. Full CD spectra collected before and after thermal melting are overlaid, with the thermal melt at 222 nm shown on the right.

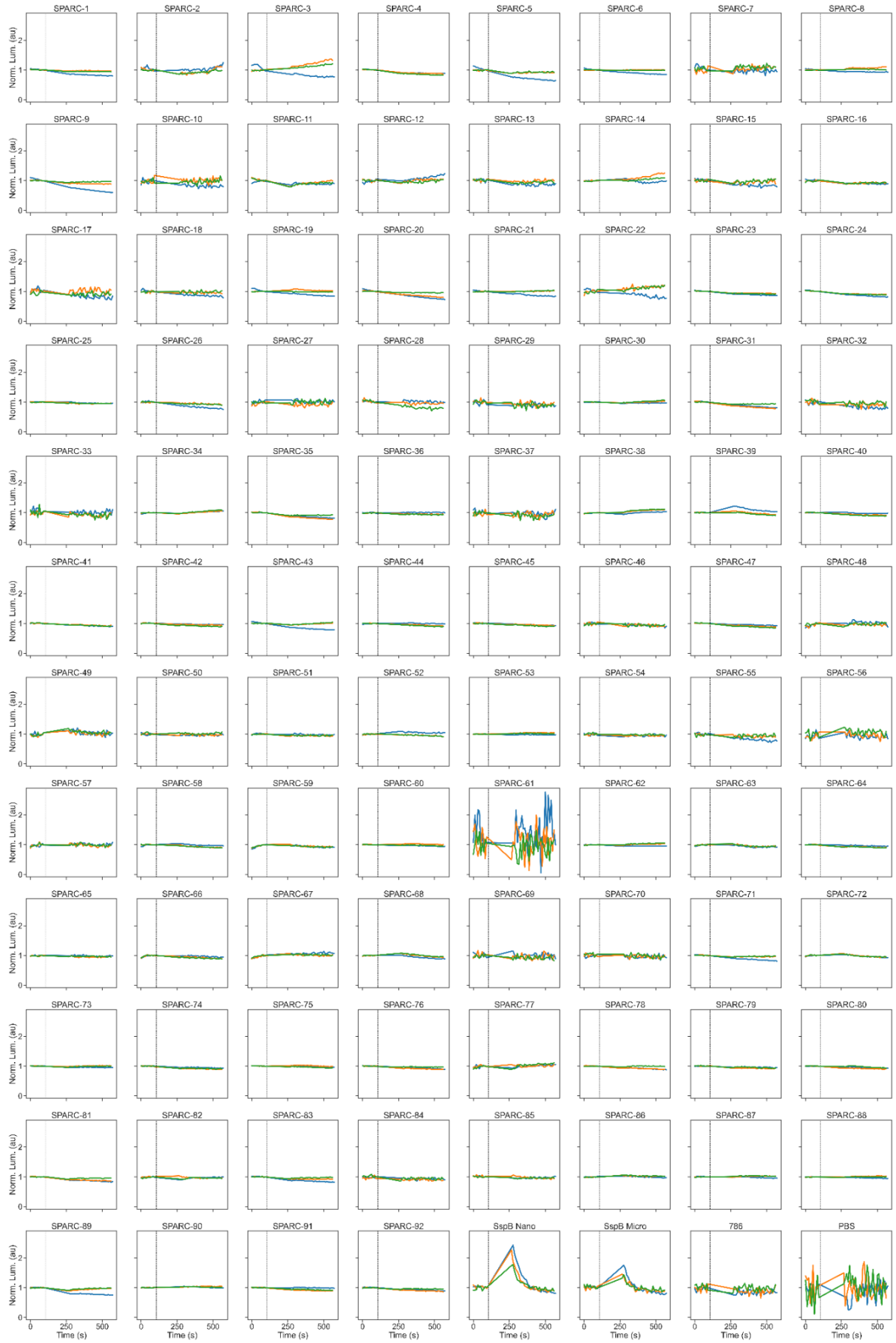

**Supplementary Fig. 6: Split-luciferase data for SPARC designs in lysate.** Three consecutive illuminations (blue, orange, green) are shown as overlays. Illumination time point is indicated by dashed vertical line and the luminescence is normalised to baseline recorded in the dark before first illumination. SspB-Nano and SspB-Micro were measured in parallel in an identical setup as positive controls (bottom right).

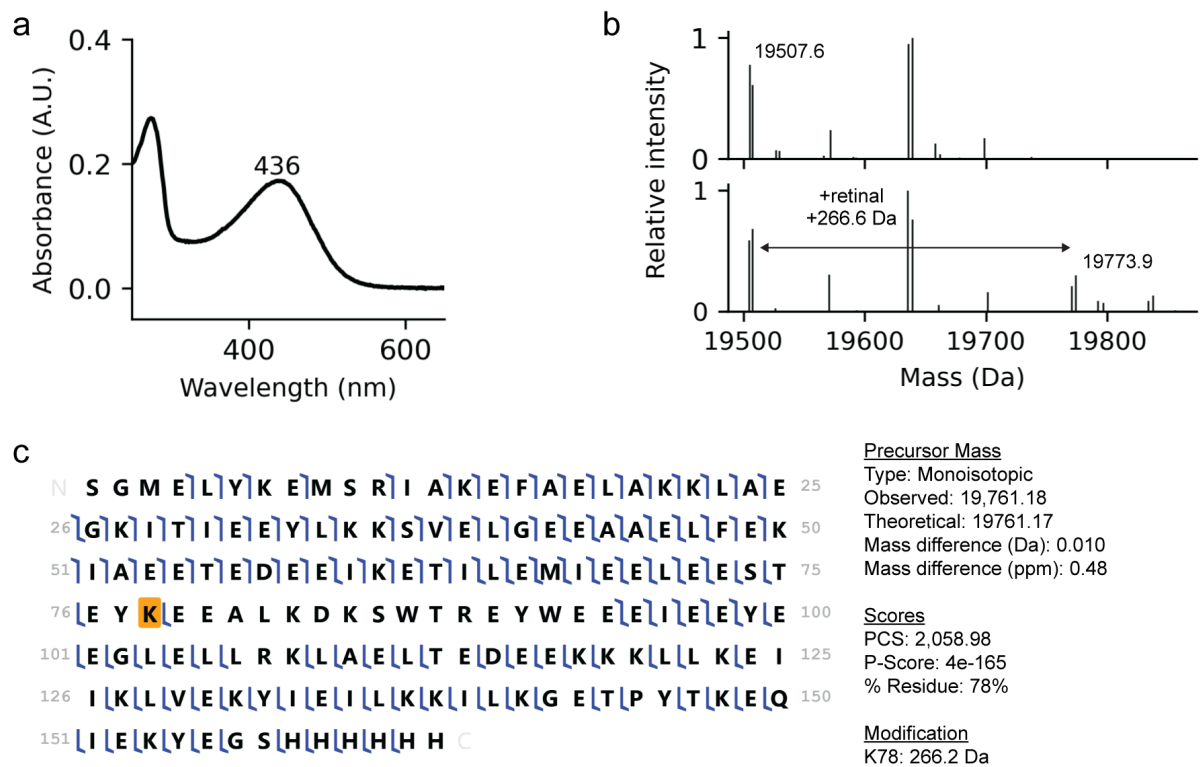

**Supplementary Fig. 7: Validation of retinal binding to SPARC-39.** **a**, Absorbance spectrum of SPARC-39 incubated with 11-*cis*-retinal. **b**, Native mass spectrum of SPARC-39 (upper) and SPARC-39 incubated with 11-*cis*-retinal. **c**, Top down MS fragmentation map for SPARC-39. A blue line in each sequence marks a detected fragment from top-down MS, with a tail towards the *N*- or *C*-terminus indicating a *b*- or *y*-fragment respectively. Location of the 11-*cis*-retinal adduct (+266.2 Da) is indicated by an orange box. % Residue indicates achieved proteoform sequence coverage.

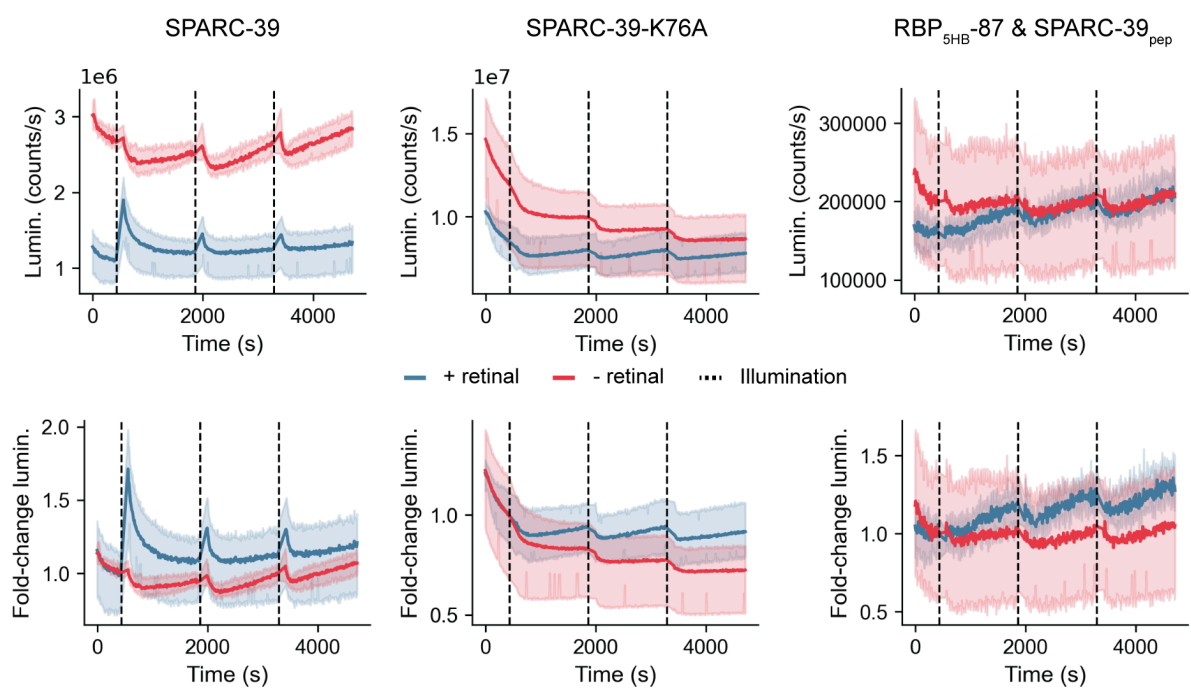

**Supplementary Fig. 8: Split luciferase data for purified SPARC-39.** K76 is the lysine residue designed for retinal Schiff base formation. RBP-87 is the parent RBP from which SPARC-39 was designed using the described multistate design pipeline. As RBP-87 has no associated peptide as part of its design it was mixed with the peptide from SPARC-39. Shaded area is the standard deviation of $n=3$ independent replicates. All samples are at 500 nM.

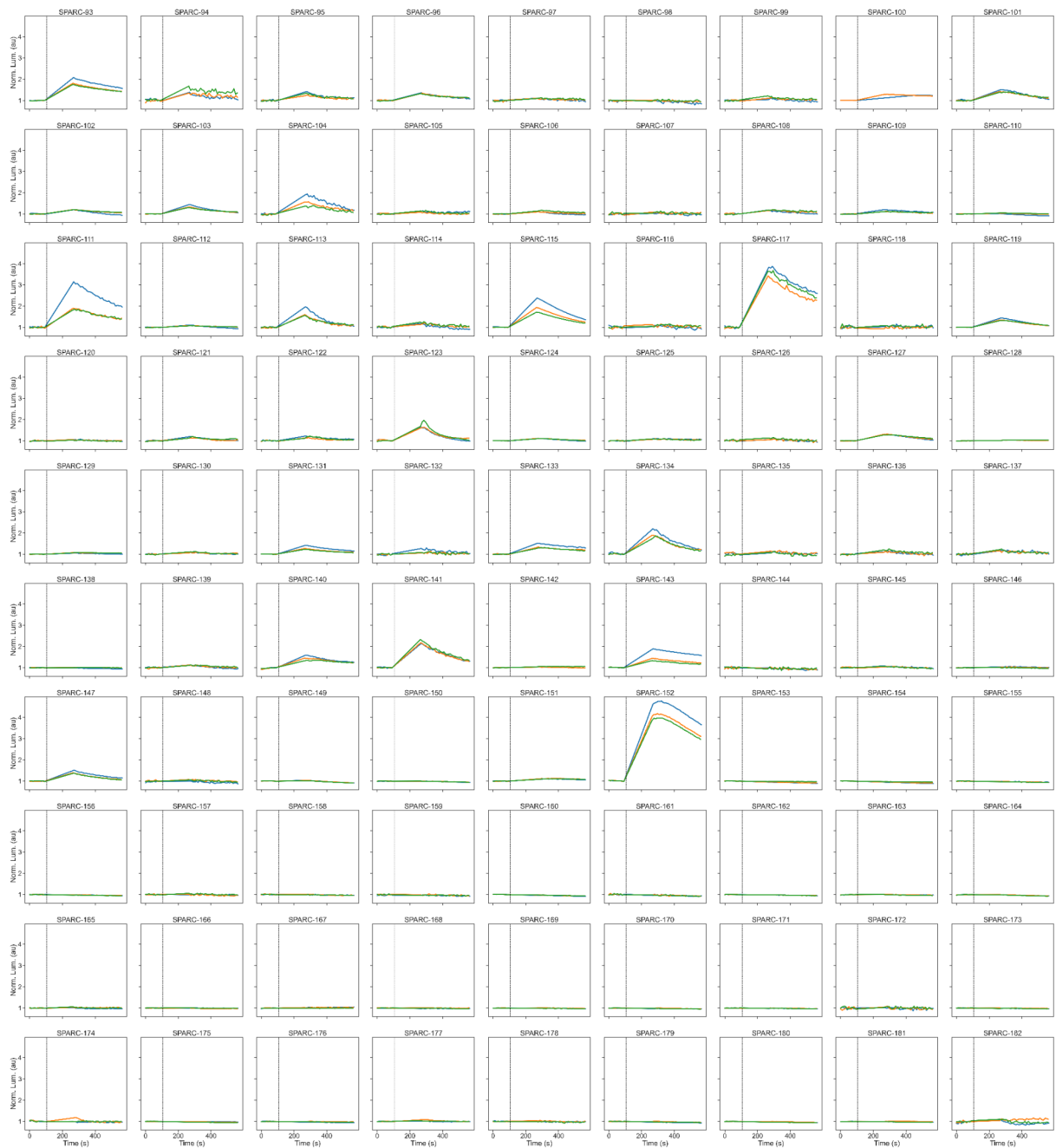

**Supplementary Fig. 9: Split-luciferase data for optimised SPARC designs in lysate.** Three consecutive illuminations (blue, orange, green) are shown as overlays. Illumination time point is indicated by dashed vertical line and the luminescence is normalised to baseline recorded in the dark before first illumination. The top 6 rows (SPARC-93 to 152) are sequence optimised using the peptide binder. The bottom 6 rows (SPARC-153 to 182) are peptide binders scaffolded into 3 or 4-helix bundles without additional sequence design.

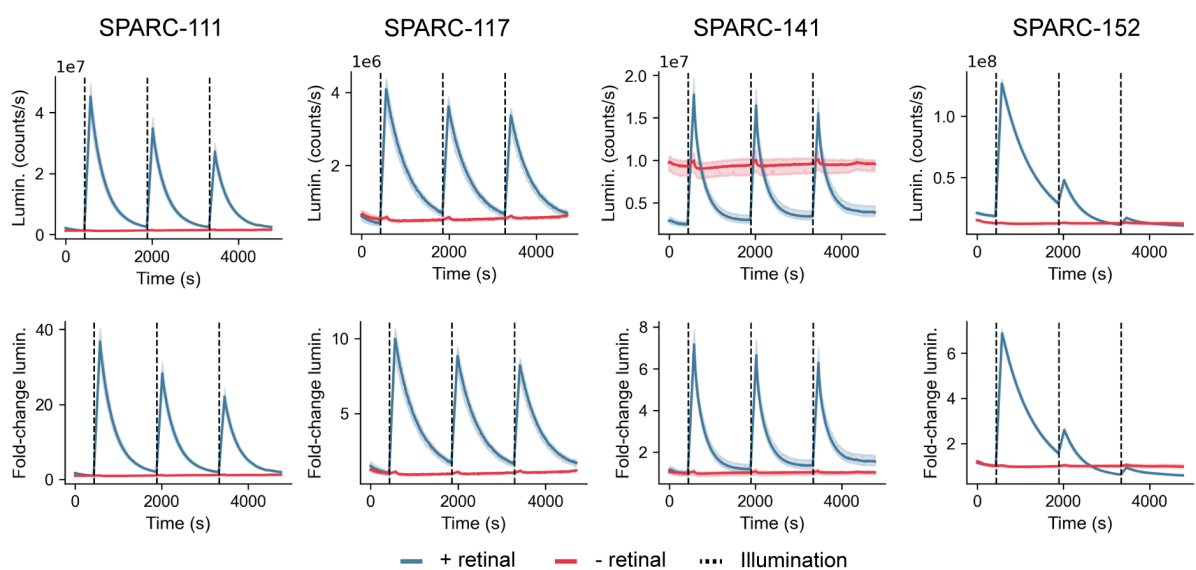

**Supplementary Fig. 10: Split-luciferase assay for purified optimised SPARC proteins.** Shaded area is the standard deviation of n=3 independent replicates. All samples are at 500 nM.

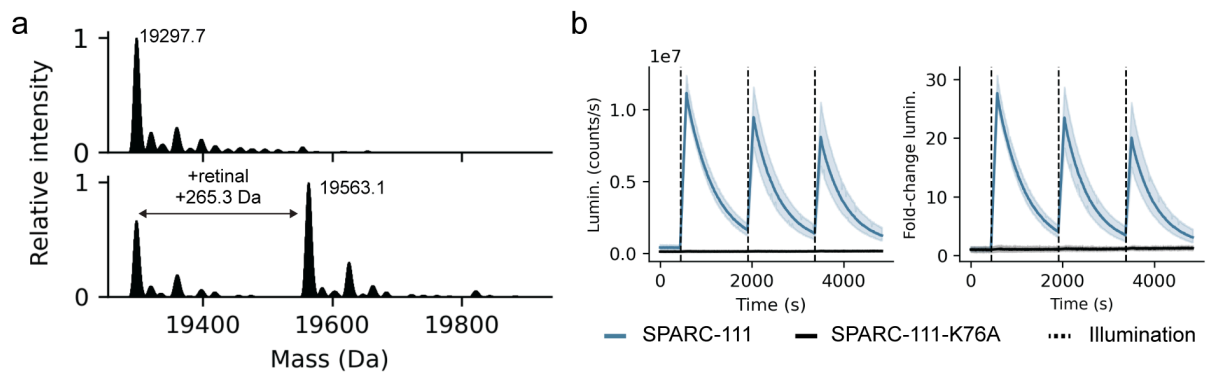

**Supplementary Fig. 11: Validation of retinal binding to SPARC-111.** **a**, Native MS of SPARC-111 and SPARC-111 incubated with 11-*cis*-retinal. **b**, Split luciferase assay of purified SPARC-111 and SPARC-111-K76A. Shaded area is the standard deviation of n=3 independent replicates. All samples are at 500 nM.

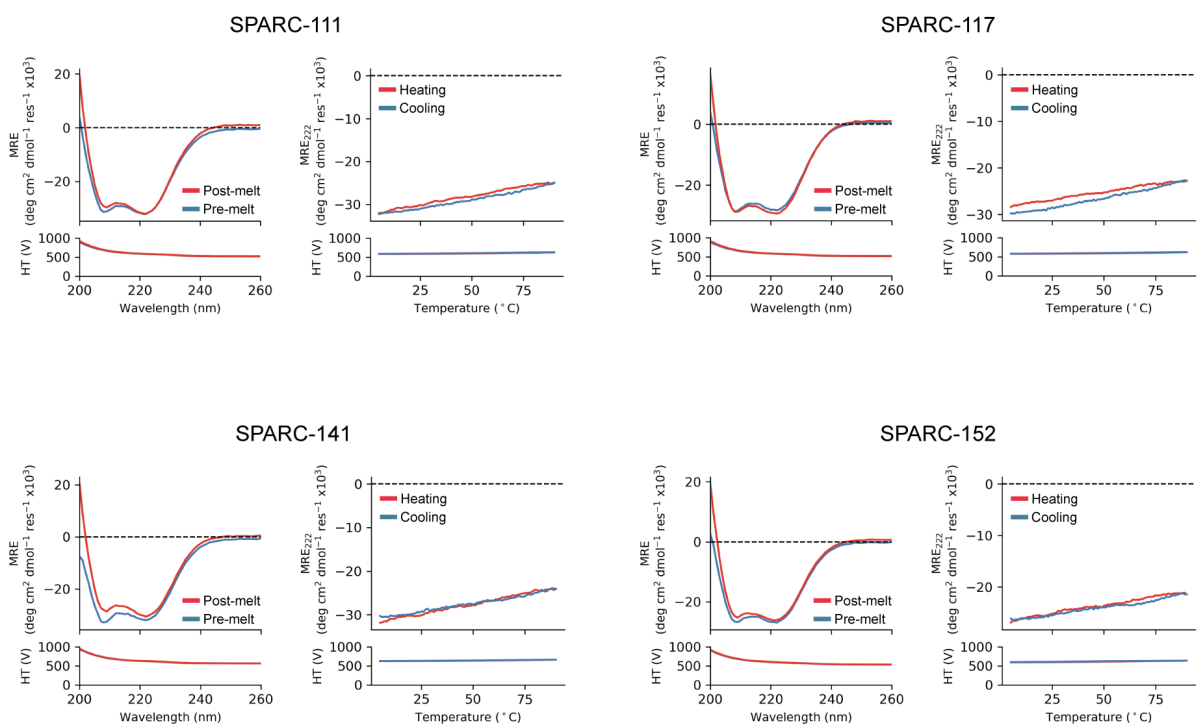

**Supplementary Fig. 12: Secondary structure and thermal stability of SPARCs.** Measurement of secondary structure and thermal stability by circular dichroism (CD) spectroscopy. Full CD spectra collected before and after thermal melting are overlaid, with the thermal melt at 222 nm shown on the right.

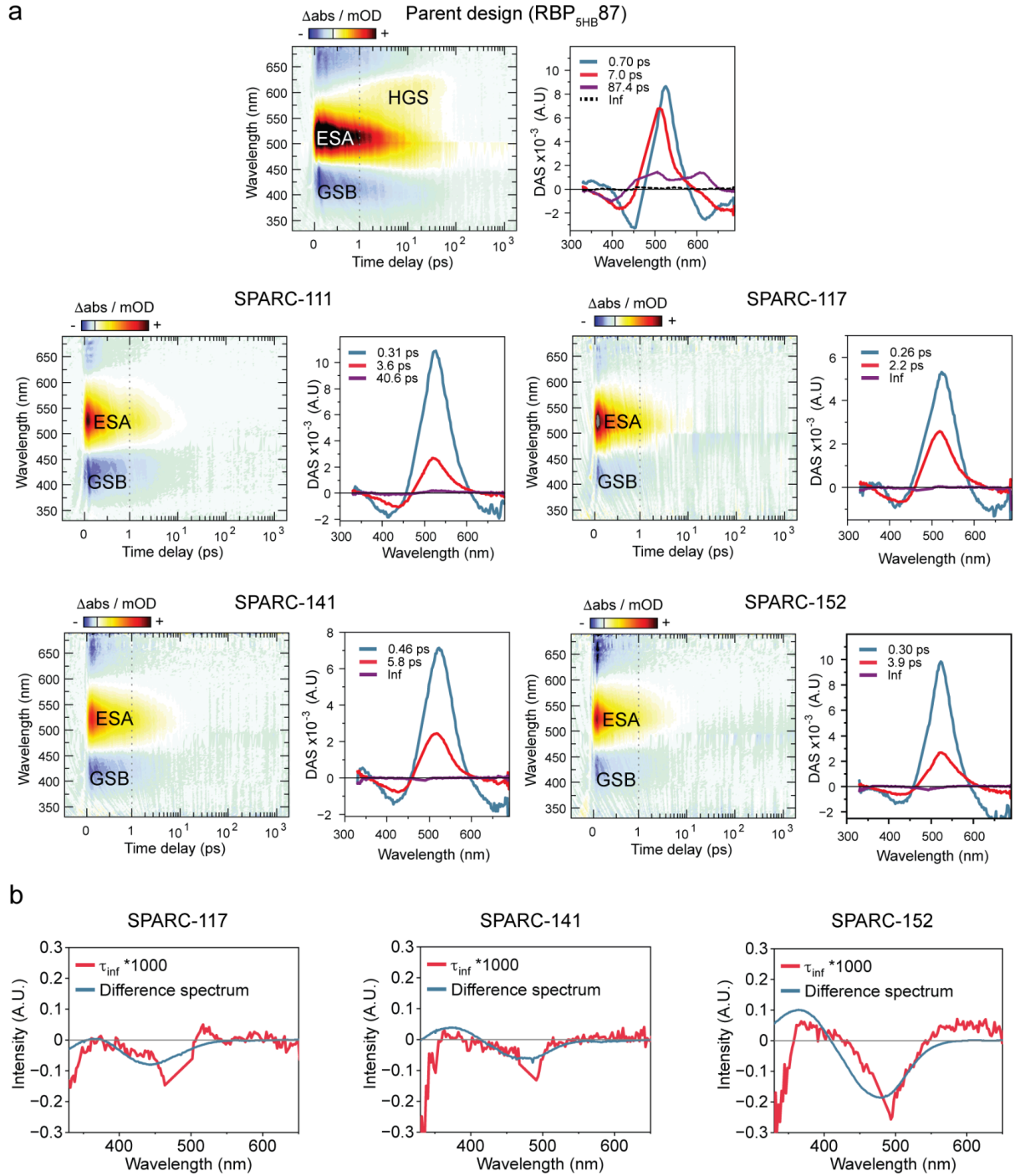

**Supplementary Fig. 13: Ultrafast spectroscopy of RBP-87 and SPARC proteins. a,** Transient absorption spectra (left) and decay associated spectra (right) resulting from a global lifetime analysis of the data. **b,** Scaled infinity lifetime ($t_{\text{inf}}$) spectra depicting the evolution at 2 ns and steady state difference spectra (after-before the measurement). The overlap shows that the remaining ground state bleach and photoproduct formation do not recover.

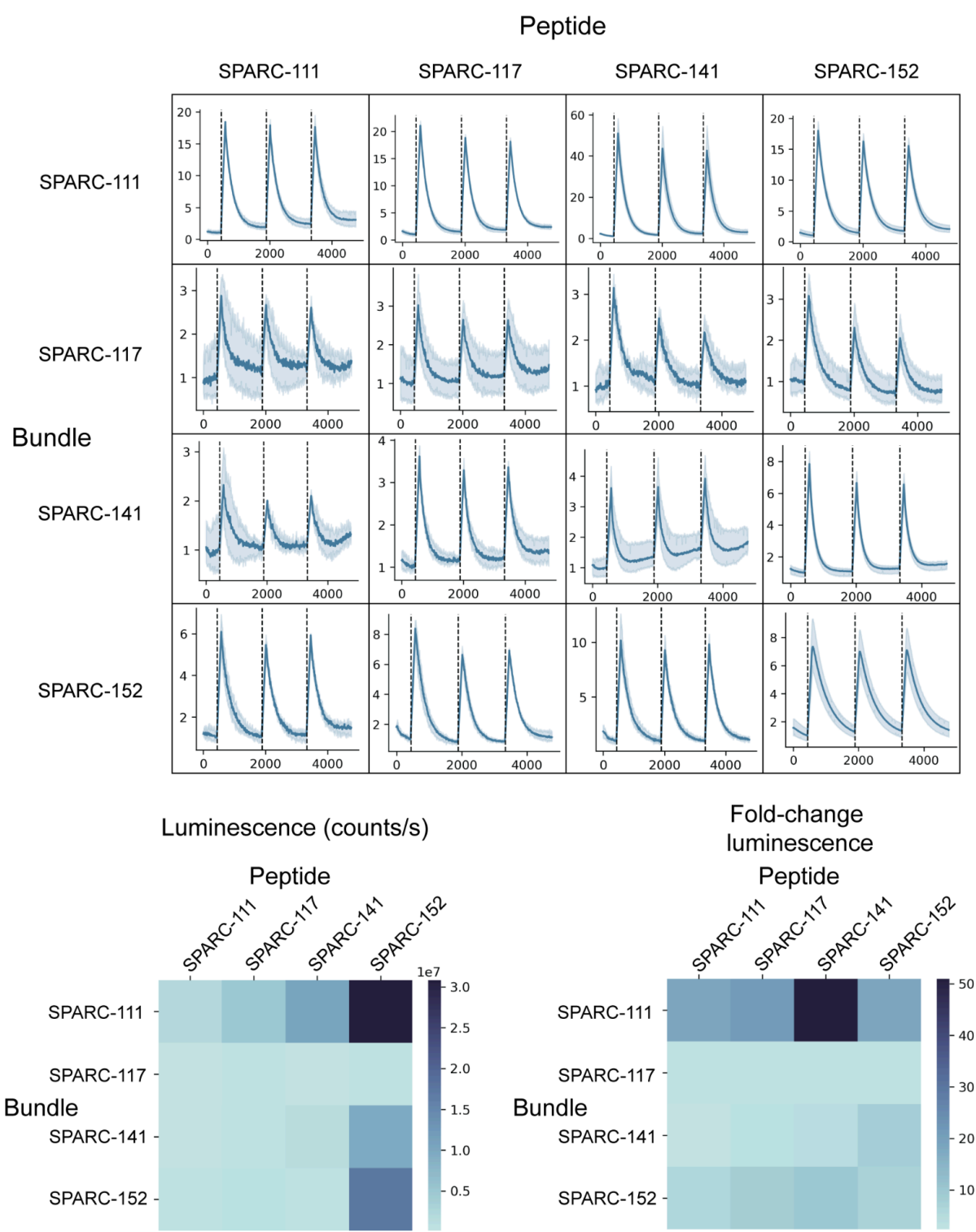

**Supplementary Fig. 14: Orthogonality matrix of optimised SPARC designs.** Normalised luminescence traces for each SPARC protein and SPARC peptide mixed at 100 nM (upper) in a split-luciferase assay. Shaded area is standard deviation of $n=3$ independent replicates. Recorded maximum luminescence (counts/s) for each pair, and the maximum recorded fold-change luminescence for each pair (lower).

### 5-helix bundle

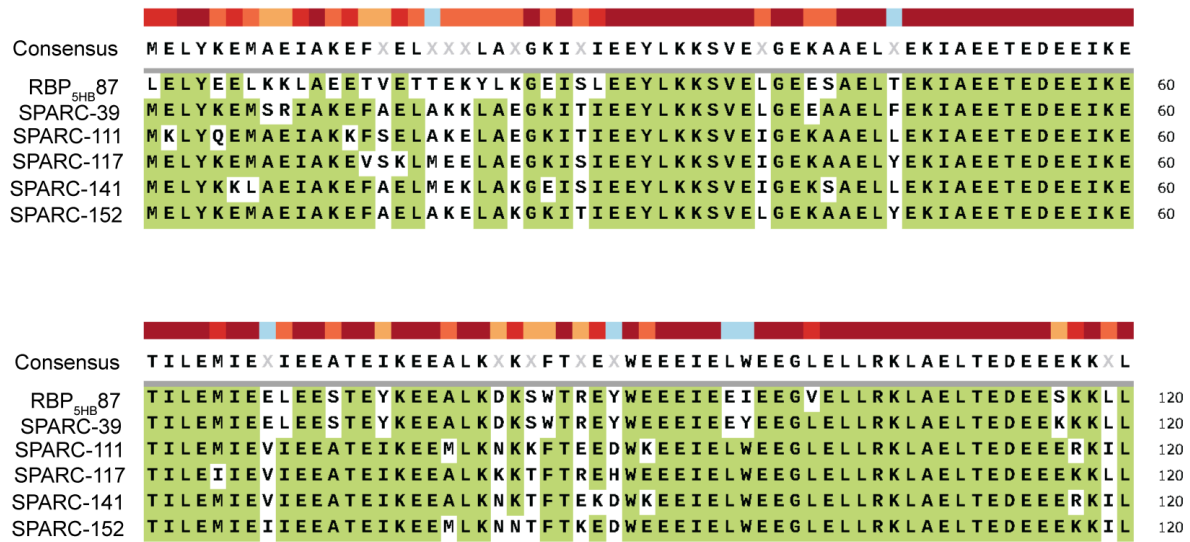

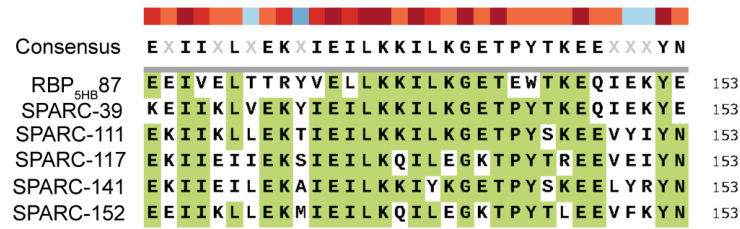

### Peptide

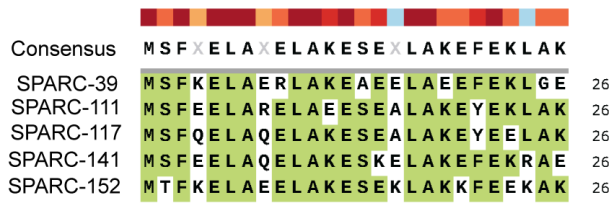

**Supplementary Fig. 15: Sequence alignment of SPARC designs.** Green highlighting indicates consensus of 50%.

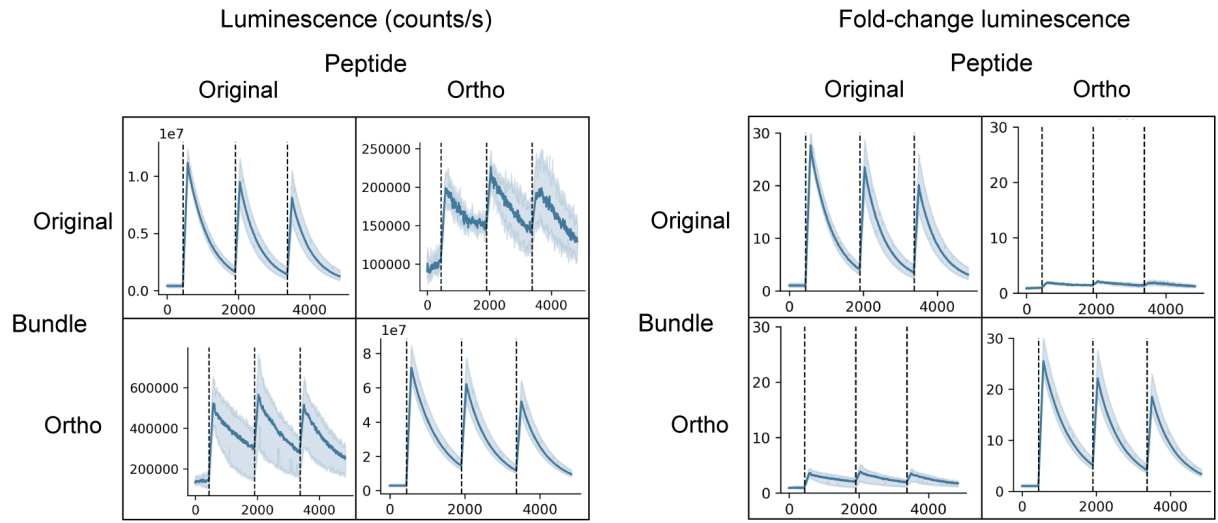

**Supplementary Fig. 16: Orthogonality matrix of charge-swapped SPARC-111.** Raw luminescence (left) and fold-change (normalised luminescence) of purified SPARC-111 (original) and SPARC-111<sub>ortho</sub> in a split-luciferase assay at 250 nM. Shaded area is standard deviation of n=3 independent replicates.

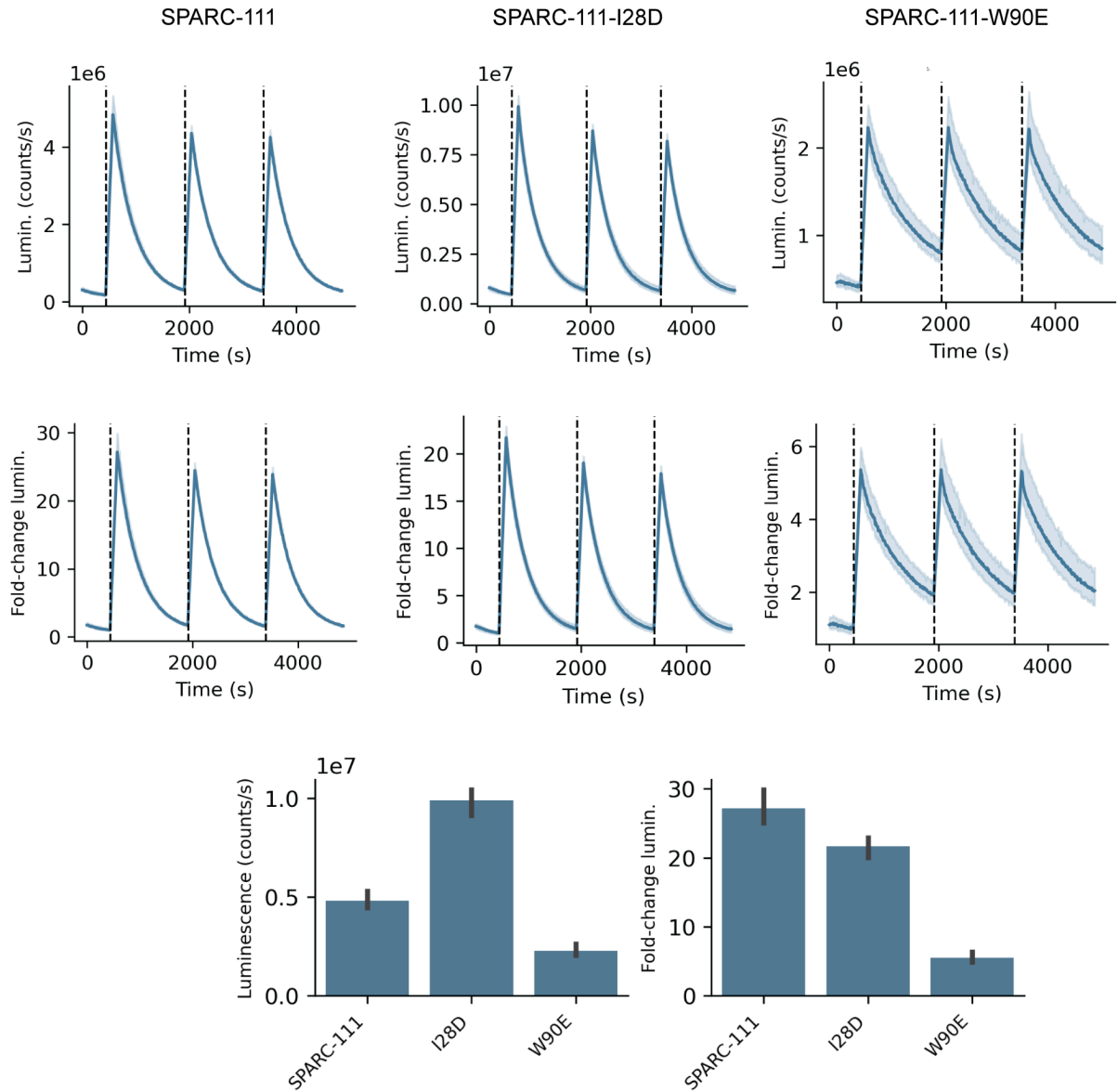

**Supplementary Fig. 17: Split luciferase assay of SPARC-111 colour mutants.** Raw and normalised luminescence of SPARC-111 and I28D and W90E point mutants. Shaded area is standard deviation of $n=3$ independent replicates at 250 nM. Below is the max recorded luminescence (counts/s) and max fold-change luminescence from each measurement.

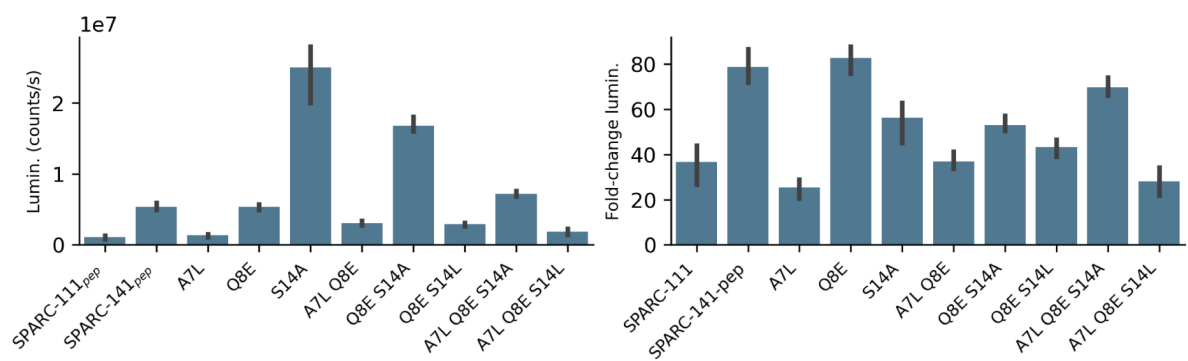

**Supplementary Fig. 18: Split luciferase assay of SPARC peptide mutants.** The max recorded luminescence (left) and max fold-change luminescence (right) from n=3 independent replicates at 100 nM.

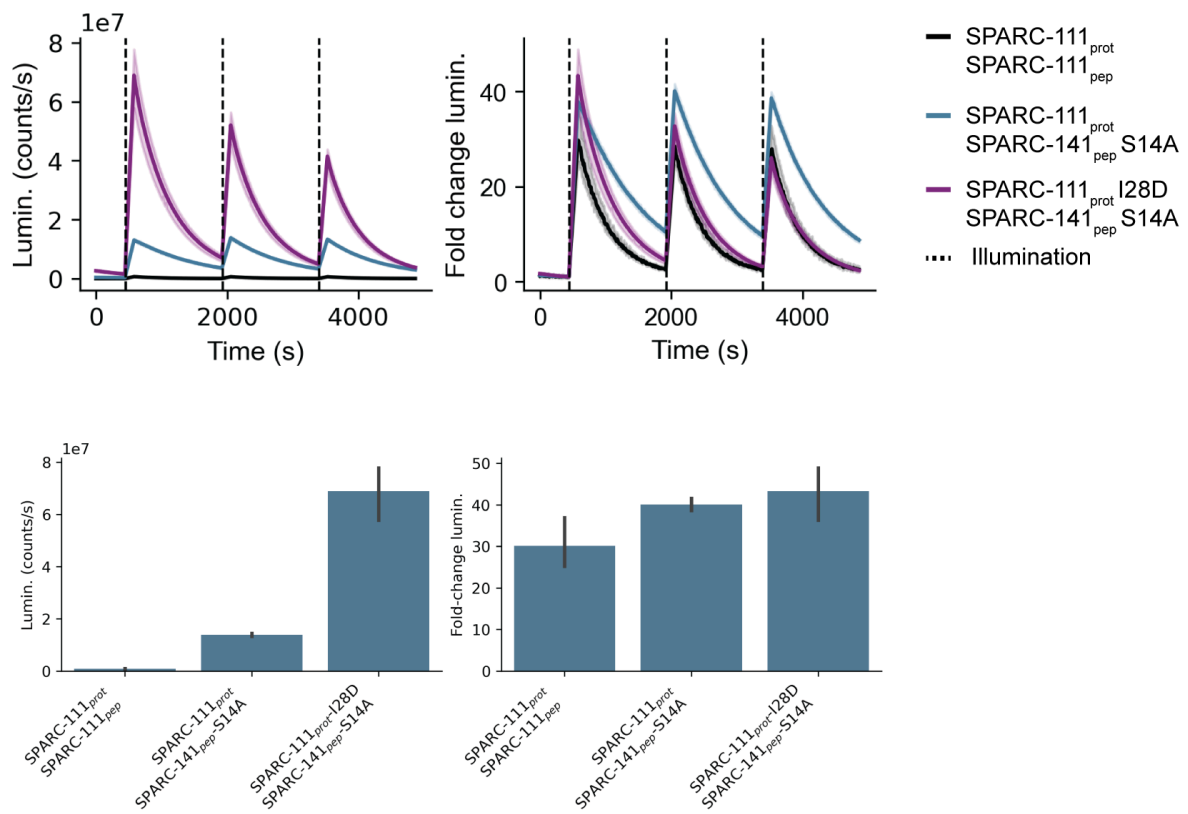

**Supplementary Fig. 19: Split luciferase assay of combined SPARC mutants.** Split luciferase traces (upper) and the max recorded luminescence (lower left) and max fold-change luminescence (lower right) from $n=3$ independent replicates at 100 nM.

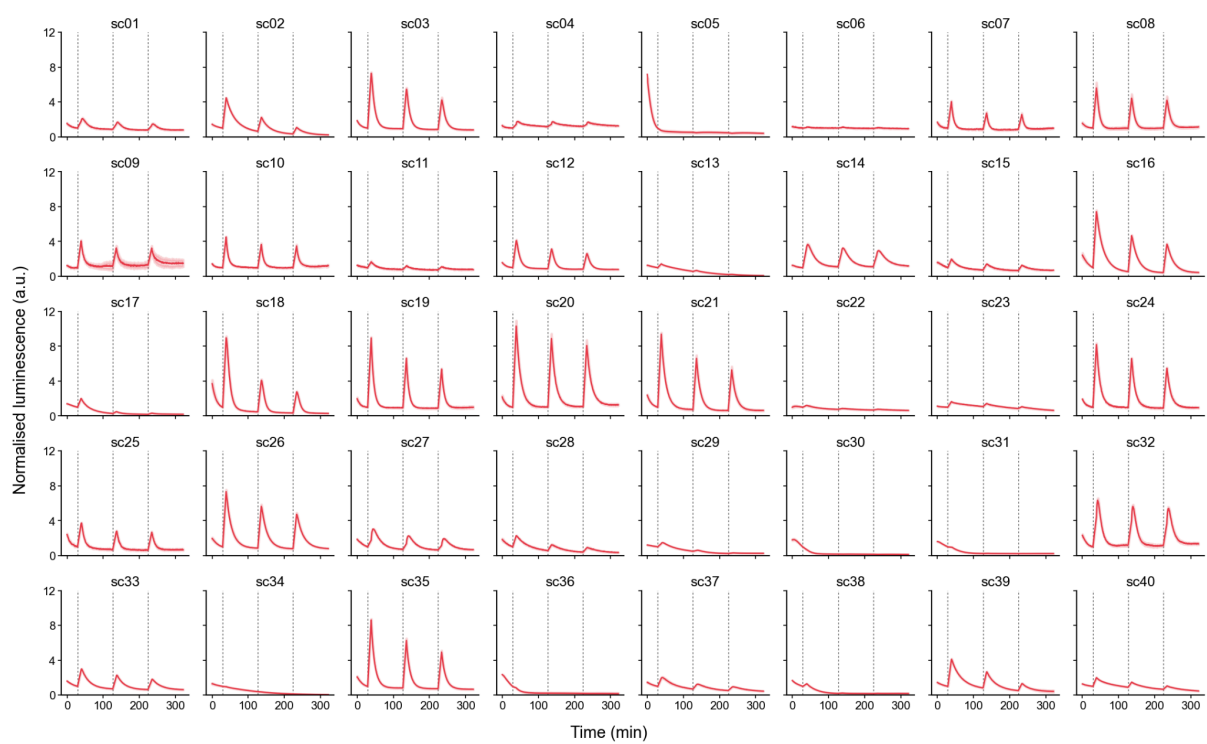

**Supplementary Fig. 20: Split-luciferase data for scaffolded SPARC peptides in lysate.** Normalised luminescence (fold-change) following three consecutive illuminations. Illumination time point is indicated by dashed vertical line and the luminescence is normalised to baseline recorded in the dark before first illumination.

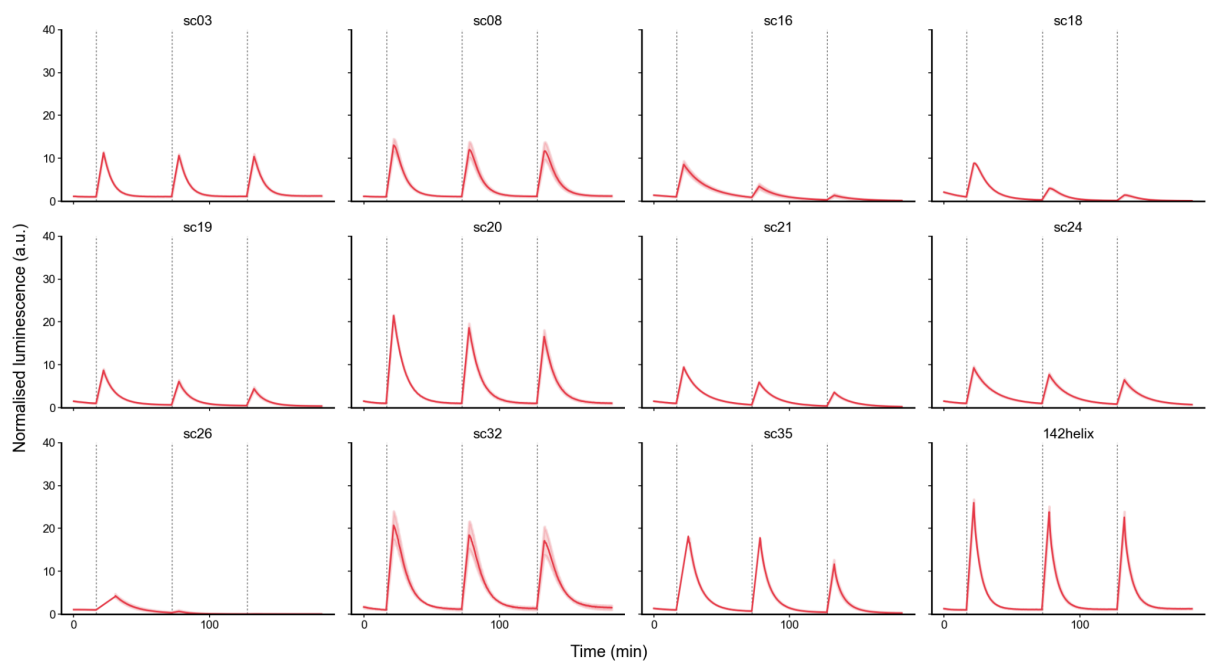

**Supplementary Fig. 21: Split-luciferase assay for purified scaffolded SPARC peptides.** Shaded area is the standard deviation of $n=3$ independent replicates. Illumination time point is indicated by dashed vertical line and the luminescence is normalised to baseline recorded in the dark before first illumination. All samples are at 250 nM.

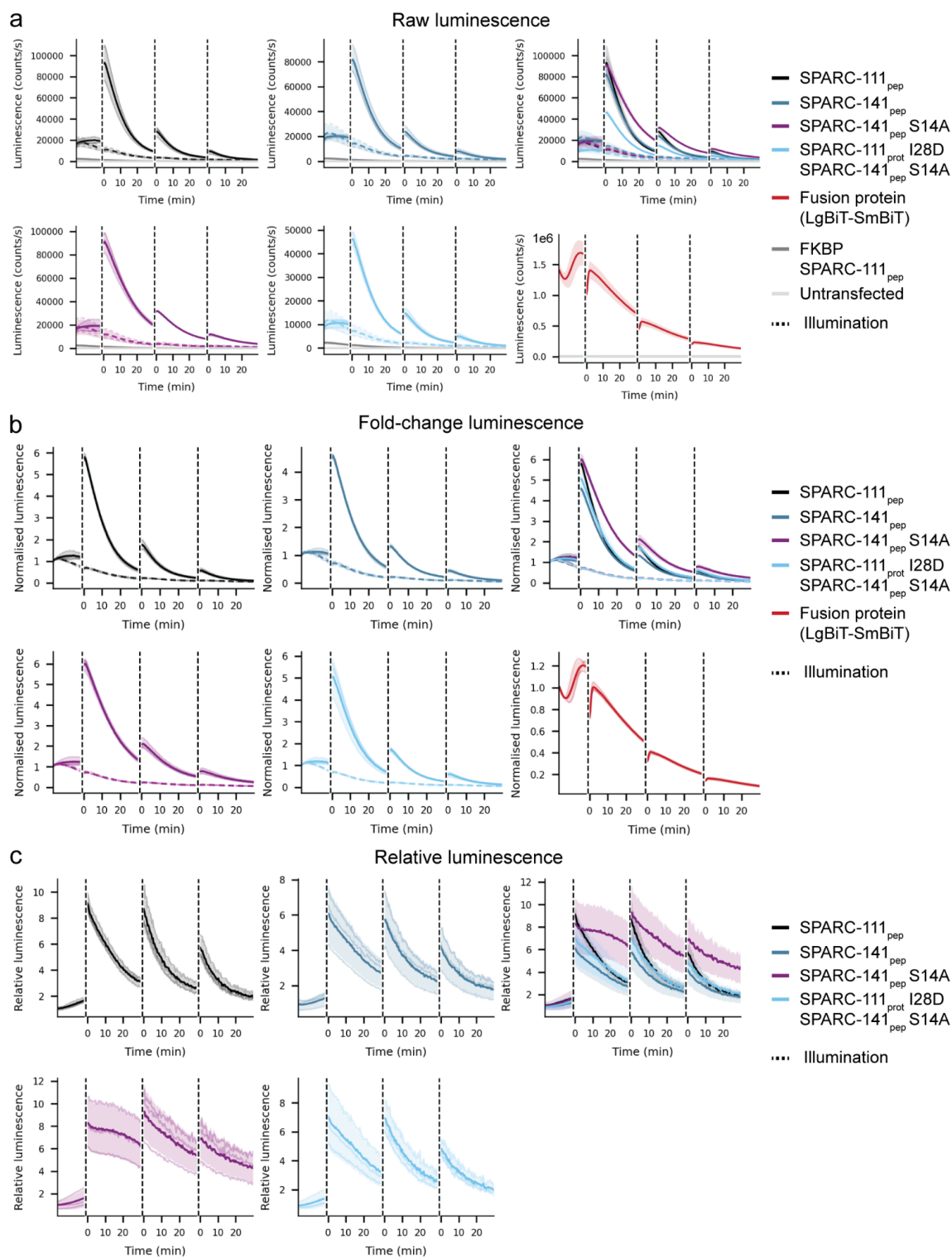

**Supplementary Fig. 22: Light-activated split-luciferase in HEK-293T.** **a**, Raw luminescence following three successive illuminations. Shaded area is the standard deviation of $n=3$ independent biological replicates. Dotted lines correspond to each construct without added 11-*cis*-retinal. FKBP was used as a split-luciferase fusion that would not be expected to interact with SPARC-111<sub>pep</sub>. **b**, Fold-change in luminescence from pre-illumination. **c**, The change in luminescence relative to the decay in luminescence for samples without 11-*cis*-retinal, to account for depletion of luciferase substrate.

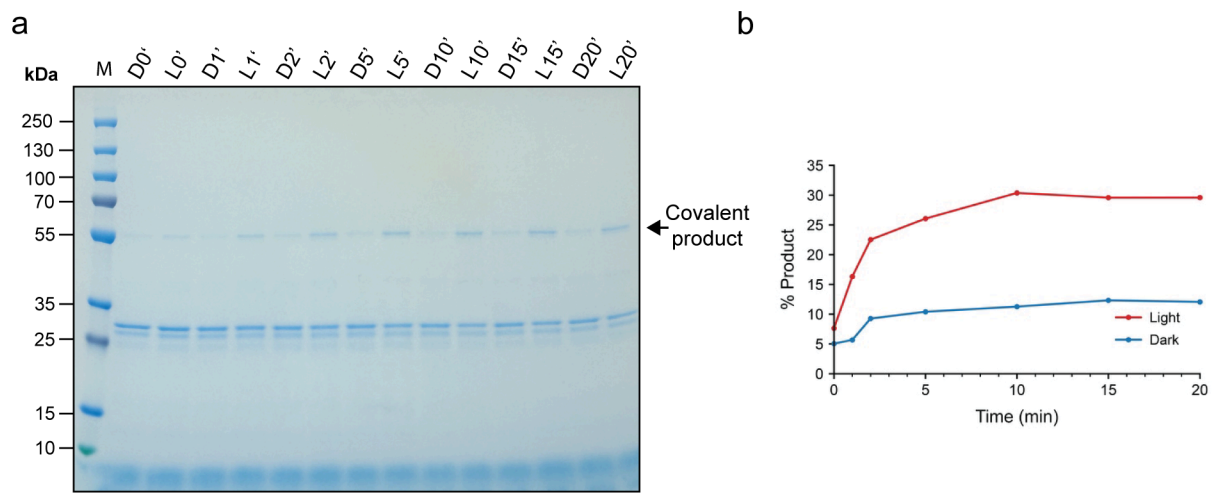

**Supplementary Fig. 23: *In vitro* SpyTag/SpyCatcher lock.** **a**, SDS-PAGE of samples illuminated (L) or kept in the dark (D) following incubation from 0 to 20 minutes. **b**, Quantification of **a**, where % product is the density of the covalent product at 55 kDa over the total density of the lane.

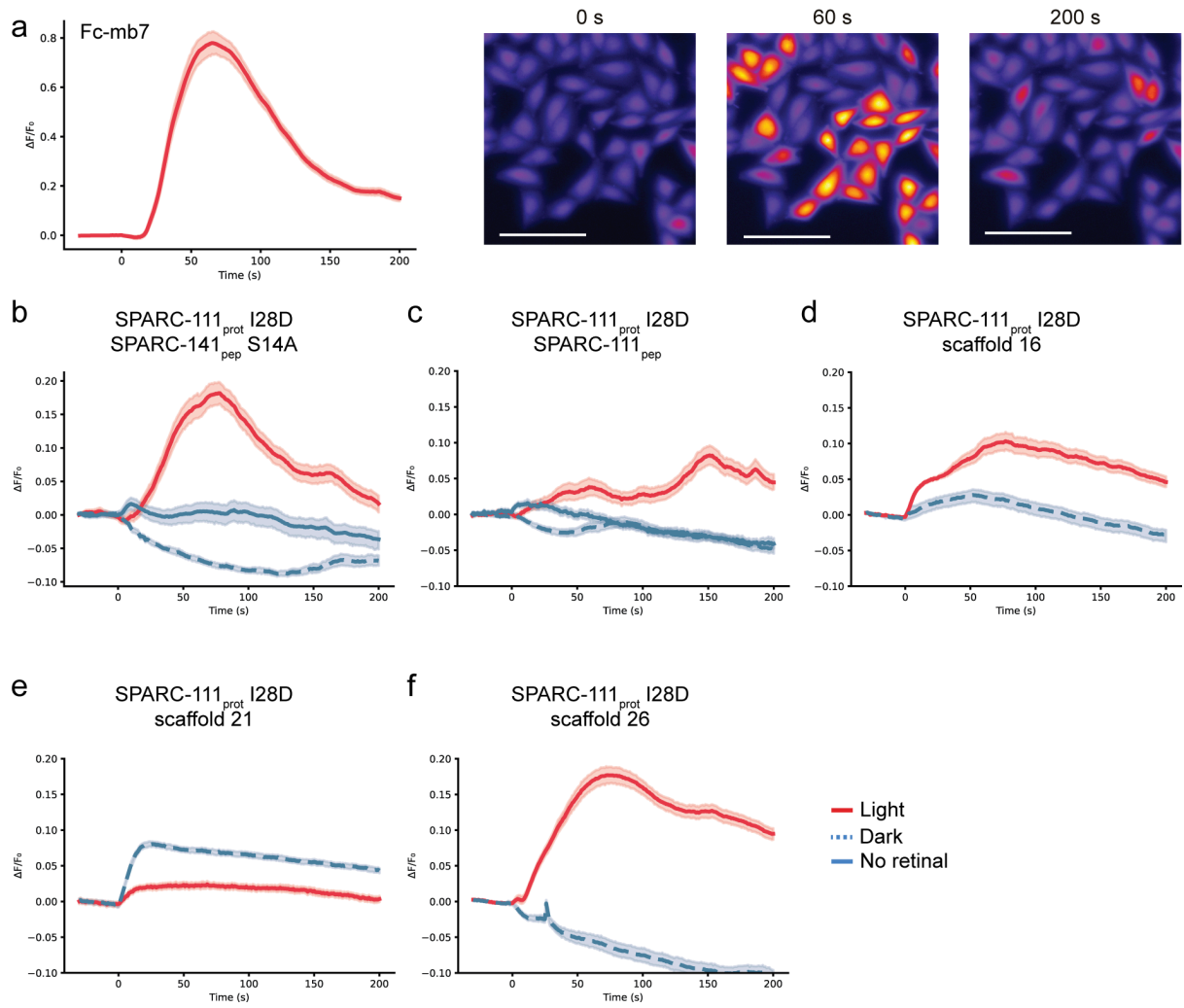

**Supplementary Fig. 24: Intracellular calcium release triggered by light-dependent FGFR1c dimerisation.** **a**, positive control (constitutive mb7 dimer) at 1 μM with representative microscopy images (scale bar 100 μm). Two biological replicates shown. **b-f**, FGFR1c activation for different SPARC pairs and scaffolded peptides. Treatment at 1 μM. **b**, 4, 1, and 4 biological replicates shown for light, dark, and no retinal condition, respectively. **c**, 2, 3, and 2 biological replicates shown for light, dark, and no retinal condition, respectively. **d-e**, 1 and 2 biological replicates shown for light and dark condition, respectively. **f**, 2 biological replicates shown for light and dark condition.

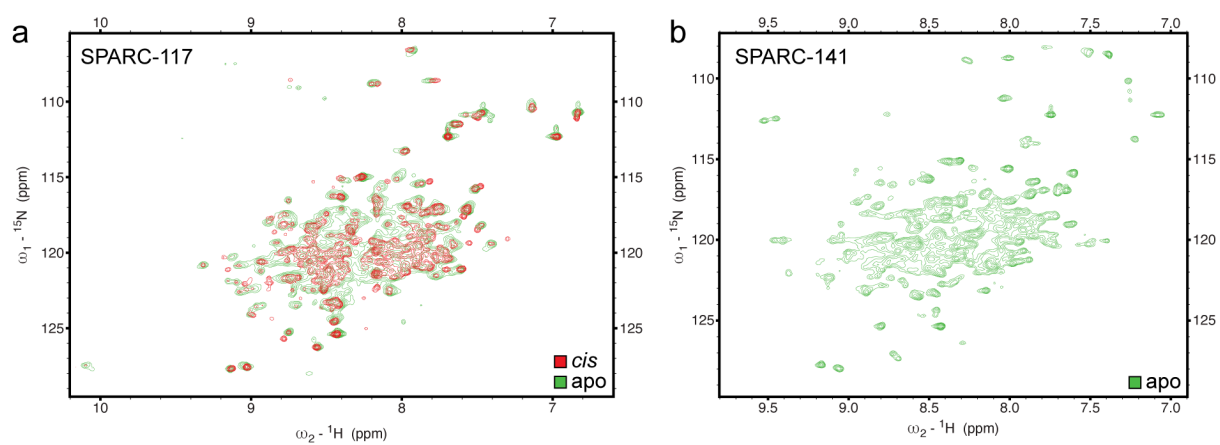

**Supplementary Fig. 25: Retinal binding by NMR.** **a**, HSQC spectra of apo (green) and holo (11-*cis*, red) SPARC-117. **b**, HSQC spectrum of apo SPARC-141.

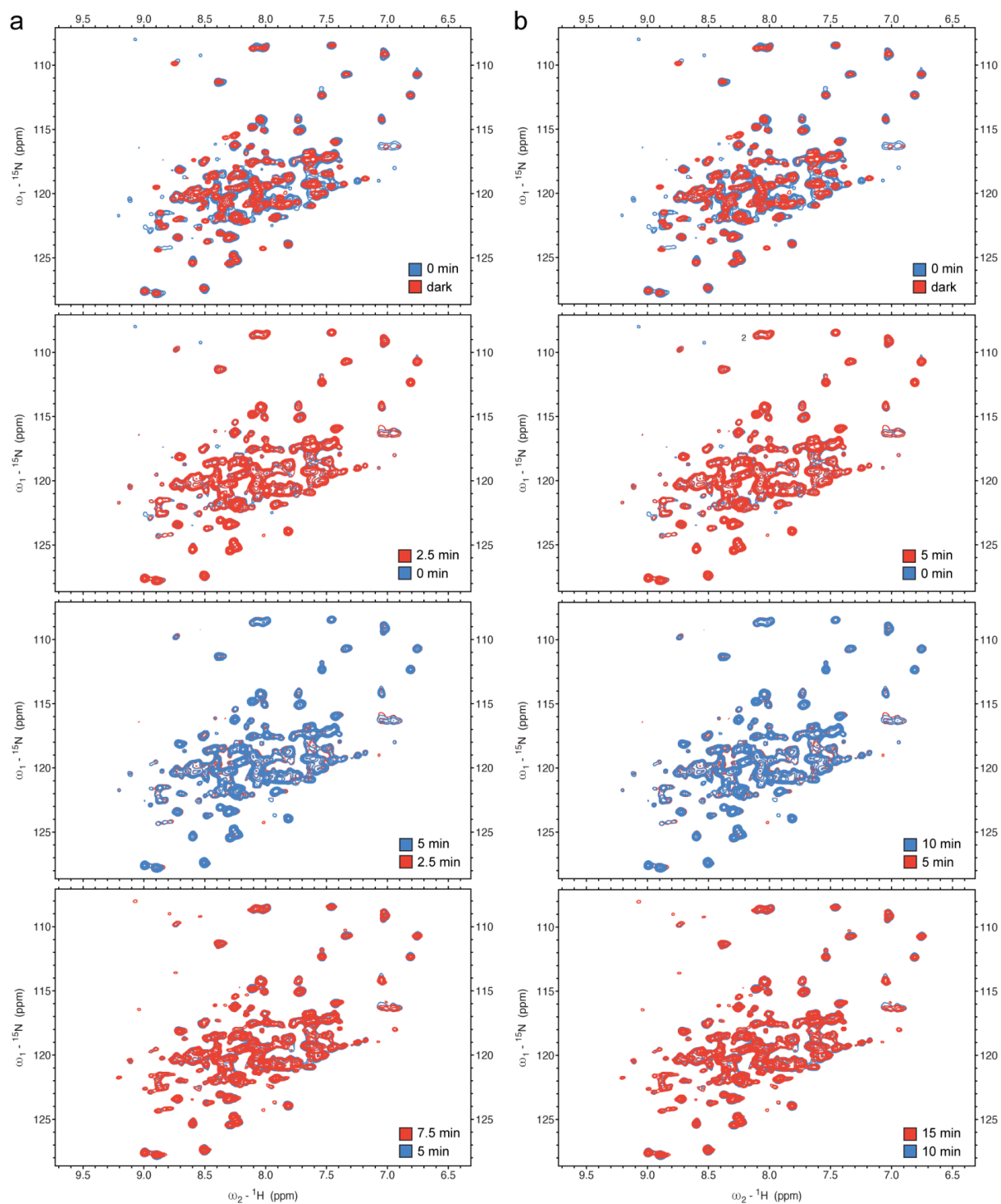

**Supplementary Fig. 26: Time series after *in situ* illumination of SPARC-111.** **a**, Series of SOFAST-HMQC $^1\text{H}$, $^{15}\text{N}$ spectra of SPARC-111<sub>prot</sub> loaded with 11-*cis* retinal before and after illumination *in situ*. **b**, Series of SOFAST-HMQC $^1\text{H}$, $^{15}\text{N}$ spectra of SPARC-111<sub>prot</sub> loaded with 11-*cis* retinal and in the presence of SPARC-111<sub>pep</sub> before and after illumination *in situ*.

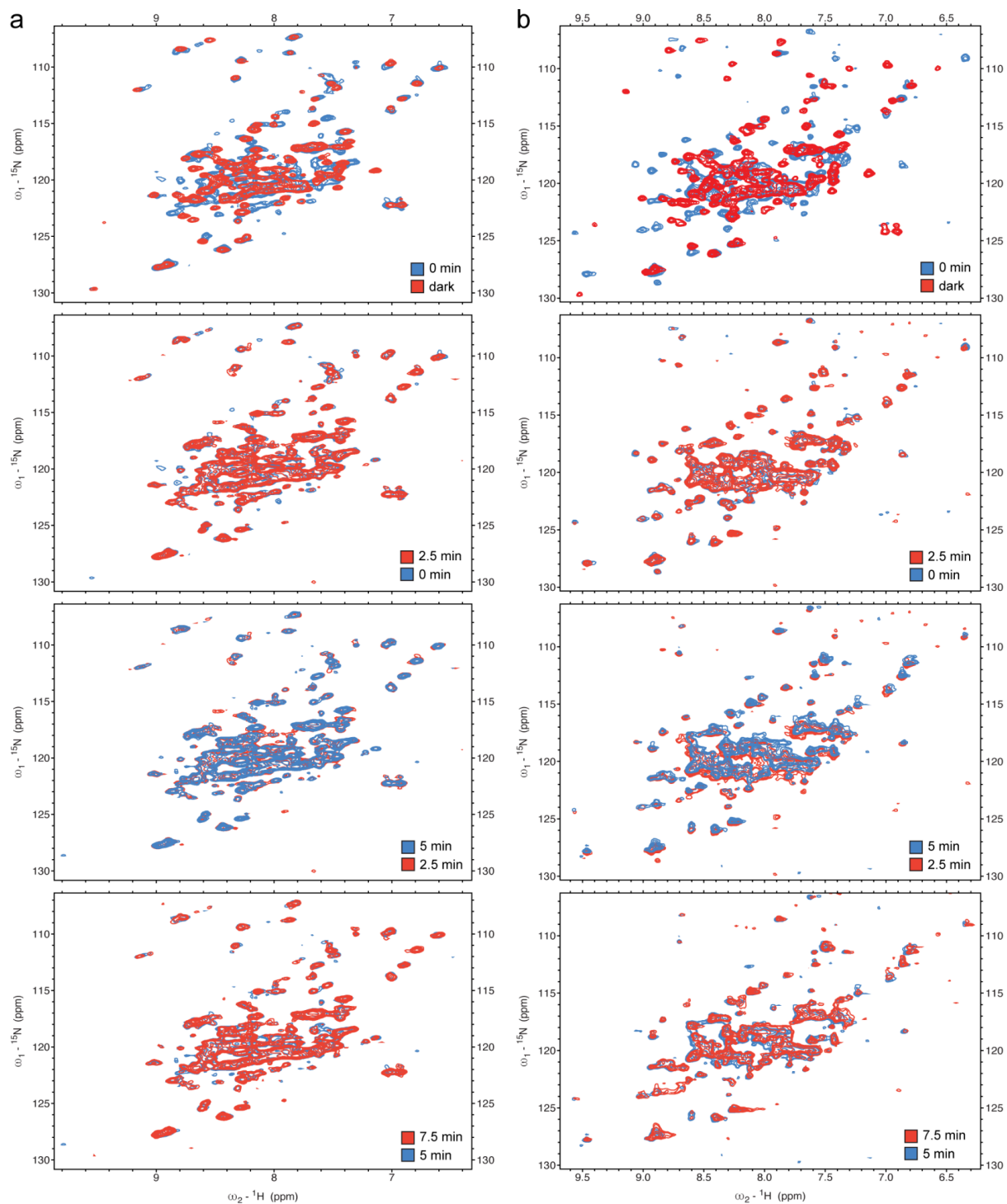

**Supplementary Fig. 27: Time series after *in situ* illumination of SPARC-152.** **a**, Series of SOFAST-HMQC $^1\text{H}$, $^{15}\text{N}$ spectra of SPARC-152<sub>prot</sub> loaded with 11-*cis* retinal before and after illumination *in situ*. **b**, Series of SOFAST-HMQC $^1\text{H}$, $^{15}\text{N}$ spectra of SPARC-152<sub>prot</sub> loaded with 11-*cis* retinal and in the presence of SPARC-152<sub>pep</sub> before and after illumination *in situ*.

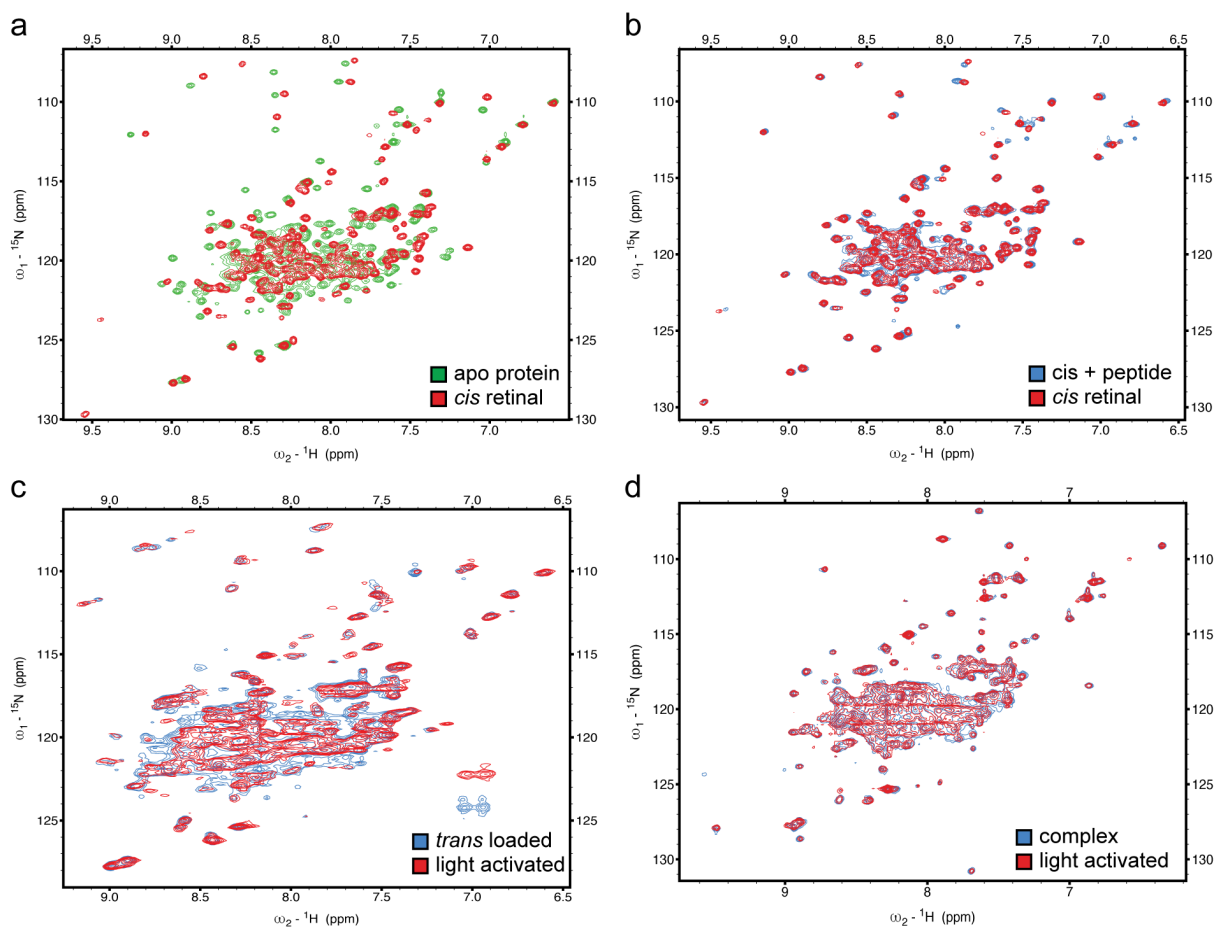

**Supplementary Fig. 28: NMR experiments on SPARC-152, including *in situ* illumination.** **a**, Overlay of apo and holo (11-*cis*-retinal) HSQC spectra. **b**, Comparison of HSQC spectra of holo (11-*cis*-retinal) protein alone and in presence of the binding partner. **c**, Comparison of light-activated intermediate generated by shining light *in situ* (red) and by loading the protein upfront with all-*trans* retinal (blue) in absence of the binding partner. HMQC spectra were acquired in 10 min each, all with 128 increments in the $^{15}\text{N}$ dimension. **d**, Comparison of the complex generated by loading SPARC-152 with all-*trans* retinal and mixing with the binding partner (blue) and *in situ* light activation of 11-*cis*-retinal loaded SPARC-152 in presence of the binding partner (red). In this case HSQC spectra are shown.

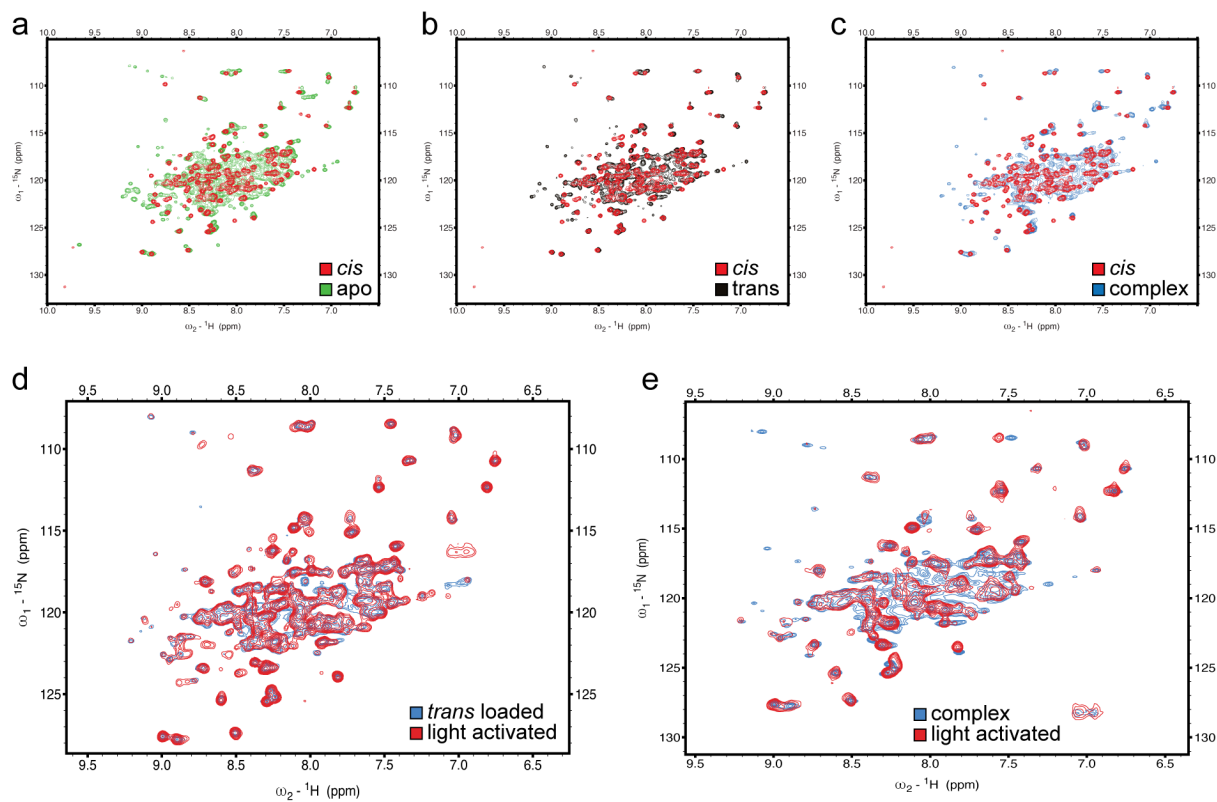

**Supplementary Fig. 29: NMR of SPARC-111.** **a**, Overlay of apo and 11-*cis* retinal loaded protein. **b**, Comparison of 11-*cis* and all-*trans* loaded protein. **c**, Comparison of 11-*cis* loaded and all-*trans* loaded protein in presence of binding partner. **d**, Overlay of all-*trans* loaded protein and *in situ* light-activated protein in absence of the binding partner. **e**, Comparison of all-*trans*-retinal-loaded protein mixed with binding partner and *in situ* light illuminated protein in presence of the binding partner. Panels **a** – **c** are HSQC spectra, **d** and **e** are HMQC spectra with 2.5 min acquisition time.

**Supplementary Fig. 30: Assay of SPARC-111 and SPARC-152 with all-*trans*-retinal.** Split luciferase assay of SPARC-111 and SPARC-152 incubated with 11-*cis* or all-*trans*-retinal.

| SPARC-117 (PDB ID: 34XO) |  |
| --- | --- |
| **Data collection** |  |
| Space group | P 1 21 1 |
| Cell dimensions |  |
| *a*, *b*, *c* (Å) | 35.79, 82.47, 55.25 |
| $\alpha$, $\beta$, $\gamma$ (°) | 90.00, 102.60, 90.00 |
| Resolution (Å) | 53.92 – 1.70 (1.73 - 1.70) |
| *R*<sub>sym</sub> or *R*<sub>merge</sub> | 0.125 (3.427) |
| number of observations | 192736 (9496) |
| *unique* | 34429 (1699) |
| *I* / $\sigma$ *I* | 7.1 (0.9) |
| CC (½) | 0.996 (0.491) |
| Completeness (%) | 99.2 (98.8) |
| Redundancy | 5.6 (5.6) |
| **Refinement** |  |
| Resolution (Å) | 53.92 – 1.70 |
| No. reflections | 33663 |
| *R*<sub>work</sub> / *R*<sub>free</sub> | 0.1900/0.2120 |
| No. atoms |  |
| Protein | 4914 |
| Ligand/ion | 114 IPE / 56 ACT / 2 MG |
| Water | 57 |
| *B*-factors |  |
| Protein | 34.7 |
| Ligand/ion | 58.9 IPE / 57.2 ACT / 42.6 MG |
| Water | 43.6 |
| R.m.s. deviations |  |
| Bond lengths (Å) | 0.0150 |
| Bond angles (°) | 2.4940 |
| Ramachandran |  |
| favoured (%) | 99.7 |
| allowed (%) | 0.3 |
| Rotamer outliers (%) | 2.6 |
| Clash score | 3.7 |

**Supplementary Table 1: Data collection and refinement statistics (molecular replacement).** The dataset was collected from a single crystal. Values in parentheses are for the highest-resolution shell.

| Spectrum | Number of crosspeaks | <sup>1</sup>H linewidths (Hz) |
| --- | --- | --- |
| SPARC-152<sub>prot</sub> apo | 259 | 48 ±52 |
| SPARC-152<sub>prot</sub> 11-*cis*-retinal | 170 | 20 ±16 |
| SPARC-152<sub>prot</sub> all-*trans*-retinal | 359 | 43 ±41 |
| SPARC-152<sub>prot</sub> all-*trans*-retinal<br>SPARC-152<sub>pep</sub> | 223 | 49 ±38 |

**Supplementary Table 2:** Numbers of crosspeaks and distribution of linewidths observed in <sup>1</sup>H,<sup>15</sup>N HSQC spectra of SPARC-152 in its 4 main forms: apo, 11-*cis*-retinal, all-*trans*-retinal without cognate peptide, and all-*trans*-retinal in presence of cognate peptide. Crosspeaks were picked automatically with the software Sparky-NMRFam after setting contours right above the noise level. Linewidths are along the <sup>1</sup>H dimension as determined by fitting all crosspeaks identified using Sparky's peak integration tool. As a reference for the number of crosspeaks, one would expect 171 accounting for asparagine and glutamine sidechains and a single proline, assuming all peptide HN correlations are observable including those from residues at the termini. All data at 308K in 20 mM MES pH 6.5, 100 mM NaCl.
